# Pronounced early differentiation underlies zebra finch gonadal germ cell development

**DOI:** 10.1101/2023.12.30.572255

**Authors:** Matthew T. Biegler, Kirubel Belay, Wei Wang, Christina Szi, Paul Collier, Ji-Dung Luo, Bettina Haase, Gregory L. Gedman, Asha V. Sidhu, Elijah Harter, Carlos Rivera-López, Kwame Amoako-Boadu, Olivier Fedrigo, Hagen U. Tilgner, Thomas Carroll, Erich D. Jarvis, Anna L. Keyte

## Abstract

The diversity of germ cell developmental strategies has been well documented across many vertebrate clades. However, much of our understanding of avian primordial germ cell (PGC) specification and differentiation has derived from only one species, the chicken (*Gallus gallus*). Of the three major classes of birds, chickens belong to Galloanserae, representing less than 4% of species, while nearly 95% of extant bird species belong to Neoaves. This represents a significant gap in our knowledge of germ cell development across avian species, hampering efforts to adapt genome editing and reproductive technologies developed in chicken to other birds. We therefore applied single-cell RNA sequencing to investigate inter-species differences in germ cell development between chicken and zebra finch (*Taeniopygia castanotis*), a Neoaves songbird species and a common model of vocal learning. Analysis of early embryonic male and female gonads revealed the presence of two distinct early germ cell types in zebra finch and only one in chicken. Both germ cell types expressed zebra finch Germline Restricted Chromosome (GRC) genes, present only in songbirds among birds. One of the zebra finch germ cell types expressed the canonical PGC markers, as did chicken, but with expression differences in several signaling pathways and biological processes. The second zebra finch germ cell cluster was marked by proliferation and fate determination markers, indicating beginning of differentiation. Notably, these two zebra finch germ cell populations were present in both male and female zebra finch gonads as early as HH25. Using additional chicken developmental stages, similar germ cell heterogeneity was identified in the more developed gonads of females, but not males. Overall, our study demonstrates a substantial heterochrony in zebra finch germ cell development compared to chicken, indicating a richer diversity of avian germ cell developmental strategies than previously known.

## Introduction

Birds have been foundational model organisms in disciplines as varied as ecology, evolutionary biology, developmental biology and neuroscience. However, compared to other groups of model organisms, the development of genetically modified avian models, including transgenic animal lines, has been quite limited. Genome editing has been most successful in the chicken (Gallus gallus), particularly through germline transmission using cultured primordial germ cells (PGCs) (Ballantyne et al., 2021b; Choi et al., 2010; Kim et al., 2010; Lavoir et al., 2006; Lyall et al., 2011; Motono et al., 2008). PGCs are early germline stem cells that give rise to egg and sperm cells. During embryonic development in birds and some reptiles, PGCs migrate from the germinal crescent to the gonadal ridges via the vascular system (Fujimoto et al., 1979; Swift, 1914). Upon reaching the developing gonad, PGCs undergo clonal expansion and apoptotic pruning before entering a quiescent state in embryonic males or committing to a meiotic fate in embryonic females (Ballantyne et al., 2021a; Cantú and Laird, 2017; Ichikawa and Horiuchi, 2023). Genome editing methods in chicken take advantage of this developmental process by harvesting PGCs from embryonic blood at Hamburger-Hamilton (HH) stage 13-16 or embryonic gonads at HH28, genetically manipulating them *in vitro*, and reintroducing them into the bloodstream of host embryos when PGC migration occurs. This allows manipulated cells to colonize the gonads as they would during normal development and subsequently contribute to the next generation.

Despite the successes in chicken, PGC-mediated genome editing and germline transmission have been difficult to apply in other bird species. Chicken is the only species for which PGCs have successfully been cultured for extended periods and maintained their commitment to the germ line (van de Lavoir et al., 2006). Short-term (2-6 passages) PGC cultures have been performed for several non-chicken species, including Japanese quail (*Coturnix japonica*), duck (*Anas platyrhyncos*), and zebra finch (*Taeniopygia castanotis*, formerly *Taeniopygia guttata castanotis*) (Chen et al., 2019; Gessara et al., 2021; Imus et al., 2014; Jung et al., 2019; Park et al., 2008; Wernery et al., 2010; Yakhkeshi et al., 2017), but long-term culture methods have not been reported. Chicken is a Galloanserae bird, which diverged over 90 million years ago with Neoaves species; in comparison most Neoaves orders diverged between 65-50 million years ago (Jarvis et al., 2014). Neoaves make up 95% of the more than 10,000 living bird species. Therefore, studies of germ cell development and subsequent establishment of a Neoaves PGC culture system is more likely to be applicable to birds generally.

An additional consideration in choosing a species to capture the diversity of avian development is the presence of the germline-restricted chromosome (GRC) in songbirds (Oscine Passeriformes). Songbirds, which include the zebra finch, constitute approximately 5,000, or half of all bird species (Ericson et al., 2003). The songbird GRC is found only in germ cells, as it is eliminated from somatic cells during embryonic development (Pigozzi and Solari, 1998; Torgasheva et al., 2019). GRC genes appear to have originated from regional duplication events of the autosomes and sex chromosomes (A chromosomes), without loss of the original genes (Borodin et al., 2022). Songbird GRC genes have only begun to be identified, as the chromosome is challenging to assemble due to the high number of highly conserved and repetitive sequences (Biederman et al., 2018; Kinsella et al., 2019). From sequencing that has been completed, it is known that the genes on the zebra finch GRC are expressed in adult testes and ovaries, and many identified genes are involved in female gonad development (Kinsella et al., 2019).

In our study, we sought to identify potential molecular differences that could explain the efficacy in *in vitro* culture conditions between chicken and zebra finch gonadal PGCs, using scRNAseq data, and compared our findings to two recent reports conducted independently (Jung et al., 2021, 2023). We found that by HH28 of both sexes, there exist two populations of zebra finch germ cells (not three as found in Jung et al., 2021), but only one at the same stage in chicken. A parallel second cluster appeared in chicken by HH36, but only in females. These two populations in zebra finch differ in expression of key transcription factors and signaling pathways that play distinct roles in germ cell biology and differentiation, as well as differential expression of GRC genes.

## Results

### Zebra finch gonadal scRNAseq identifies two germ cell populations

Male (n=2) and female (n=2) zebra finch gonads were dissected and dissociated at HH28 (around 5.5 days of development; Murray et al., 2013), a stage at which avian gonadal PGCs have previously been collected for cell culture (Choi et al., 2010; Jung et al., 2019) (Figure 1–figure supplement 1A; Supplemental table 1). Samples were processed for scRNAseq using the 10x Genomics platform, and the reads mapped against a high-quality zebra finch reference assembly (21,762 gene annotations; Supplemental table 2) produced by the Vertebrate Genomes Project (GCF_003957565.221; Rhie et al., 2021). Embryo sex was validated by W chromosome gene expression (Figure 1–figure supplement 1B). This mapping and a stringent quality control pipeline were used to remove confounding artifacts commonly seen in scRNAseq analysis (Luecken and Theis, 2019) (Figure 1–figure supplement 1C-G; Supplemental table 3). A total of 8,970 cells passed quality control.

**Figure 1.**
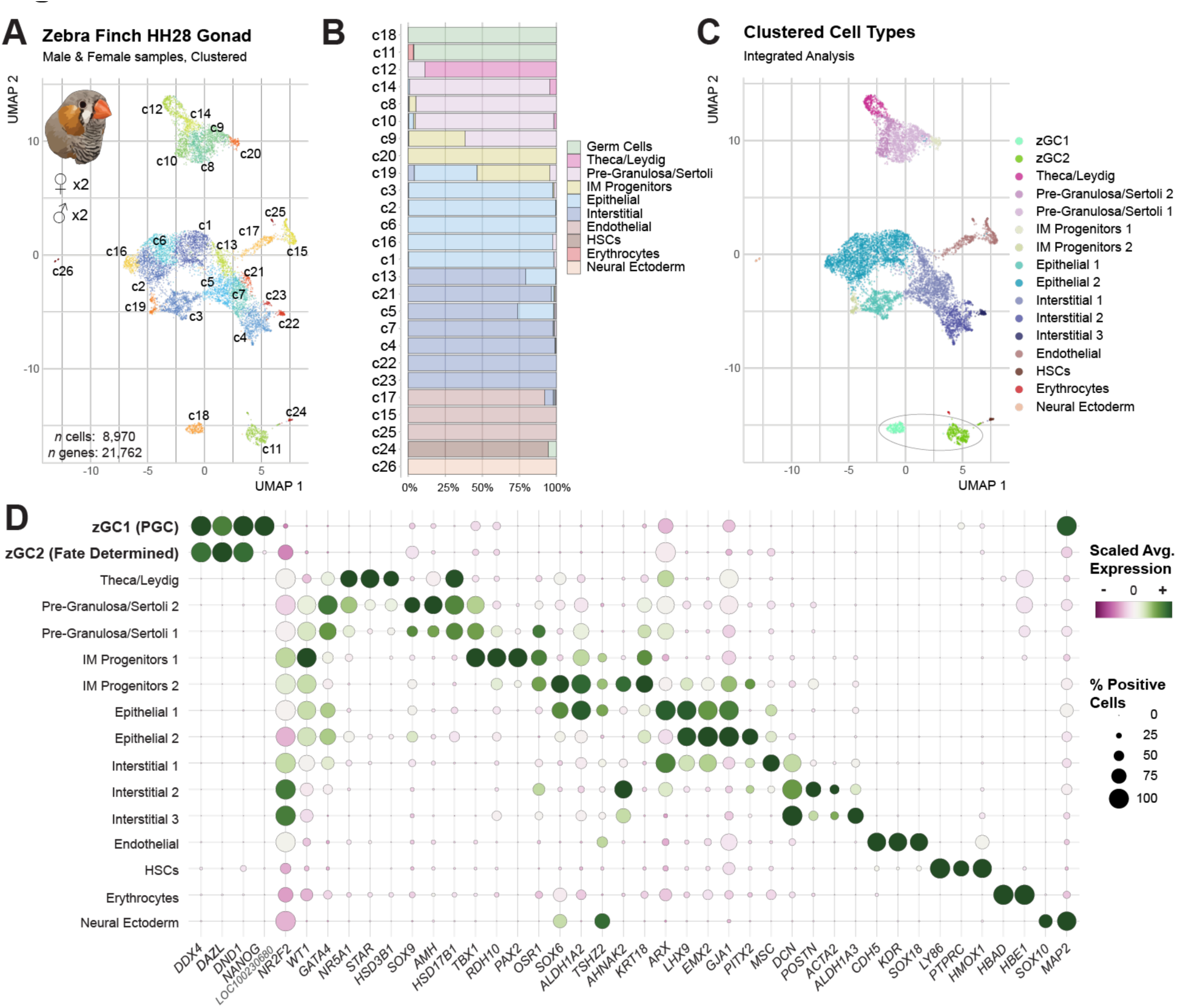
Identification of two germ cell types in the zebra finch gonad. A. UMAP plot of male and female zebra finch gonadal nearest-neighbor cell clusters at HH28. Further information on quality control and dimensional reduction for this dataset may be found in Figure 1–figure supplement 1. B. Proportional bar chart of inferred cell types present in each nearest-neighbor cluster. C. UMAP plot of male and female zebra finch clustered cell types at HH28. See Figure 1– figure supplement 2 for more information on designation. D. Dot plot of scaled expression for select gene markers of each clustered cell type.

Gene expression-based PCA analyses were visualized by UMAP dimensional reduction, with 26 nearest-neighbor clusters resolved (Figure 1A). To identify cell types among these clusters, we assign labels to a strict subset of cells marked by canonical cell type gene expression patterns (figure supplement 2A; Supplemental table 4). The gene expression profiles of these assigned cell types were then applied as a reference in a label transfer analysis (Stuart et al., 2019), inferring the cell types of the remaining cells by gene expression profile similarity (Figure 1–figure supplement 2B-D). Both male and female populations included the expected major gonadal cell types (Figure 1B) seen in other species at this stage of development (Estermann et al., 2020b; Jung et al., 2021; Stévant et al., 2019) (Figure 1–figure supplement 2B). By combining the cell-type labels with clusters, we identified several cell subtypes, including two groups of epithelial cells, three groups of interstitial cells, and two groups of putative intermediate mesodermal (IM) progenitor populations (Figure 1C).

Two distinct but hierarchically-related clusters, c18 and c11, were identified as expressing the germ cell markers *DAZL*, *DDX4* and *DND1* (Figure 1B-C; Figure 1–figure supplement 2A-B), which we broadly defined as zebra finch germ cell clusters 1 and 2 (zGC1 and zGC2). These two clusters were stably resolved across UMAPs generated with varying numbers of dimensions (Figure 1–figure supplement 1H) and nearest-neighbor clustering resolutions (Figure 1–figure supplement 1I). Both zGC clusters contained cells from males and females (Figure 1–supplemental figure 2C), indicating that clustering was not due to sex. Both clusters were also marked by increased unique molecular index (UMI) read counts and gene counts (Figure 1–figure supplement 2D), consistent with recent findings of stem cell hypertranscription (Kim et al., 2023). Interestingly, only zGC1 expressed *NANOG* (Figure 1D), a canonical marker of embryonic stem cells and PGCs (Chambers et al., 2007; Jean et al., 2015).

### The two zebra finch germ cell populations dynamically express GRC genes

We next wanted to determine the extent of expression from the GRC in the two zGC clusters. However, as the zebra finch GRC has not yet been sequenced in its entirety and no gene annotations exist in the current reference genome (GCF_003957565.2), we hypothesized that GRC gene transcripts in the zebra finch germ cells may be mismapping to conserved paralog annotations on the A chromosomes (Figure 2A). Of the high-confidence GRC candidate gene paralogs identified in a previous reference genome version (GCF_000151805.1; Kinsella et al., 2019), we identified 77 in the current reference assembly used to analyze our scRNAseq datasets (Supplemental table 5). Compared to somatic cell types in the gonad, 24 of these candidate genes were upregulated in at least one of the zGC clusters (Figure 2B and Figure 2–figure supplement 1; Supplemental table 6). These included genes related to TGF-b superfamily/SMAD signaling pathways (*BMPR1B*), RA response-mediated gene expression (*RXRA)*, and canonical PGC identity (*PRDM1*, also known as *BLIMP-1*). Additionally, 13 candidate genes were differentially expressed between the two zGC populations (log2 fold-change ≥ 0.5 and adjusted p-value ≤ 0.05). Using an aggregate of GRC candidate gene expression for UCell module analysis (Andreatta and Carmona, 2021), we saw significantly higher module scores in both zGC populations compared to somatic cell types (Figure 2C; Supplemental table 7), indicating that significant GRC gene expression was indeed being incorrectly captured as A chromosome gene expression.

**Figure 2.**
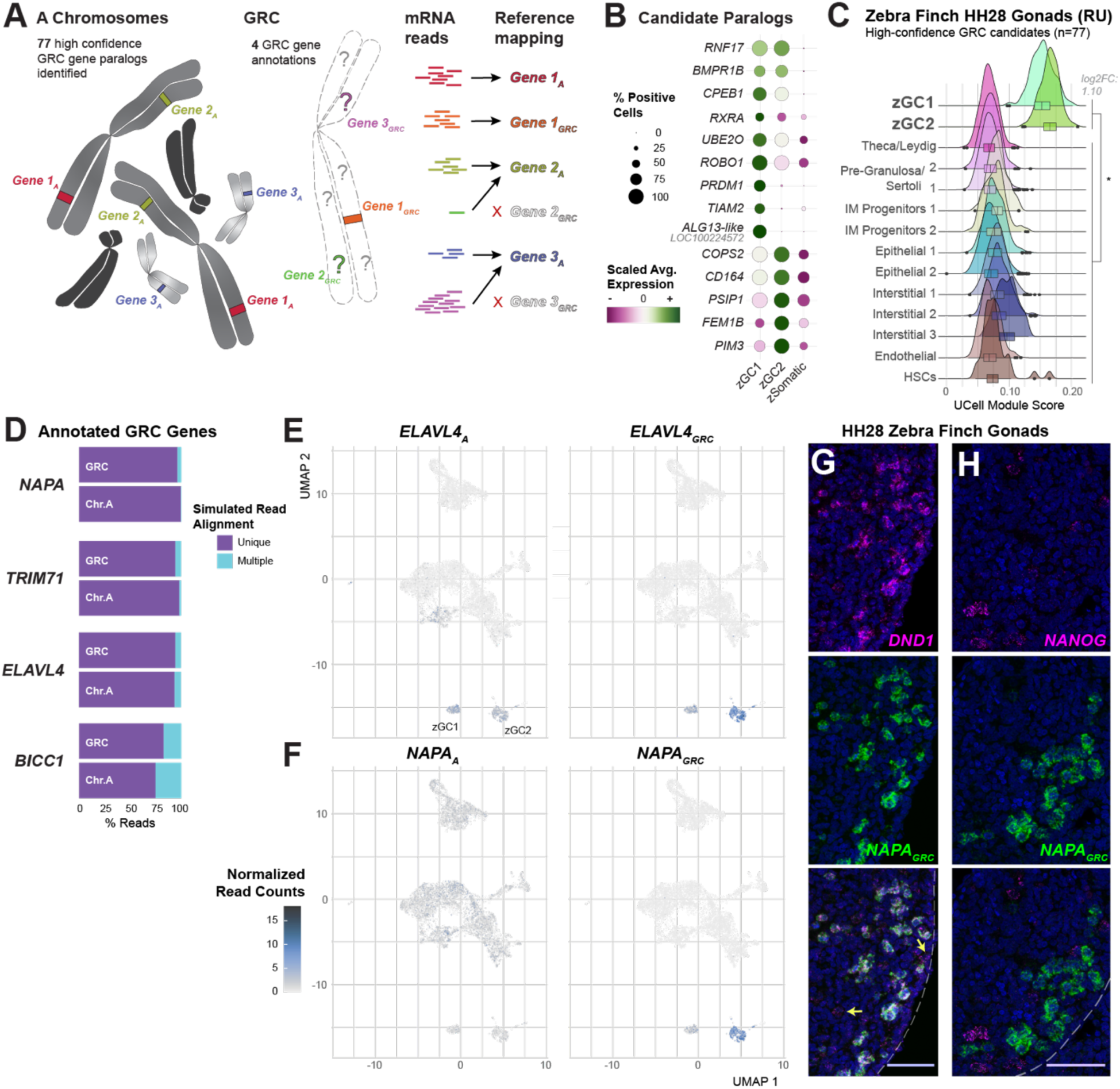
Assessment of germline-restricted chromosome genes in the zebra finch HH28 gonad. A. Diagram of putative GRC gene read mapping onto annotated somatic gene paralogs. 81 GRC candidates were identified in the current annotation used in this study, 4 of which possess available sequences for germ cell deconvolution. B. Scaled-expression dot plot of select high-confidence GRC gene candidates (Kinsella et al., 2019) between zGC and aggregate zSomatic clusters. A. Module score assessment of the 77 unmapped, high-confidence GRC gene paralogs in each clustered cell type. A Log2FC > 0.5 between zGC and zSomatic populations and a p-value ≤ 0.05 by two-sided t-test (Supplemental table 33) is denoted by *. Heatmap of expression for individual genes may be found in ED1. C. Simulated read multi-mapping assessment between GRC and A chromosome gene pairs. D. UMAP plots of zebra finch male and female HH28 zebra finch gonads overlaid with *ELAVL4A* (left) and *ELAVL4GRC* (right) gene pair expression (transcripts/10,000 UMIs) for all cell barcodes. Note the high specificity of the GRC paralog sequences with the zGC clusters, particularly in zGC2. E. UMAP plots of zebra finch male and female HH28 zebra finch gonads overlaid with *NAPAA* (left) and *NAPAGRC* (right) gene pair expression (transcripts/10,000 UMIs) for all cell barcodes. Note the high specificity of the GRC paralog sequences with the zGC clusters, particularly in zGC2. F. Dual-labeled fluorescent *in situ* hybridization of *DND1* and *NAPAGRC* in the HH28 zebra finch, showing high co-localization near the medial edge of the gonad. Arrows highlight *DND1*+ cells without *NAPAGRC* expression. Scale bar = 50µm. G. Dual-labeled fluorescent *in situ* hybridization of *NANOG* and *NAPAGRC* in the HH28 zebra finch, showing little co-localization. Scale bar = 50µm.

To further resolve the potential involvement of the GRC in zebra finch germ cell heterogeneity, four published sequences of GRC gene annotations were appended to our scRNAseq dataset: *NAPAGRC*, *TRIM71GRC*, *ELAVL4GRC* and *BICC1GRC* (Biederman et al., 2018; Kinsella et al., 2019). We quantified the extent to which the GRC gene copies map uniquely to the GRC versus the corresponding A chromosome paralogs, mapping simulated reads for each gene onto a small, simulated genome containing the eight gene annotations. We found that, on average, more than 90% of reads mapped uniquely to their respective chromosomal gene origin (Figure 2D), particularly for *TRIM71*, *NAPA*, and *ELAVL4*. This simulation demonstrated that scRNAseq reads from the closely related GRC and A chromosome paralogs can be confidently distinguished and mapped.

Mapping GRC gene expression onto the UMAP cell cluster diagram allowed us to independently verify the exclusion of the GRC from all other gonadal cell types, as expression of *NAPAGRC*, *TRIM71GRC*, *ELAVL4GRC* and *BICC1GRC* was restricted to the two germ cell clusters (Figure 2–figure supplement 2A). *TRIM71GRC* and *BICC1GRC* expression was weak compared to their respective A chromosome paralogs (Figure 2–figure supplement 2B-C), while *ELAVL4GRC* and *NAPAGRC* were expressed at higher levels in the germ cells than *ELAVL4A* (Chr. 8) and *NAPAA* (Chr. 34). These GRC paralogs were particularly upregulated in the zGC2 cluster (Figure 2E-F), indicating differential gene expression between germ cell types.

We developed *in situ* hybridization probes for a minimally conserved (81.7% identity) region of the ChrA and GRC *NAPA* paralogs, which demonstrated differential signals in the zebra finch embryonic gonad (Figure 2–figure supplement 3). We validated *NAPAGRC* expression by fluorescent *in situ* hybridization (Figure 2G), which showed robust expression in a subset of *DND1+* germ cells, and further analysis showed lower expression co-localizing in *NANOG+* cells (Figure 2H). These findings indicate that the two zebra finch germ cells clusters clearly and differentially express GRC gene paralogs during early gonadal development.

### The two zebra finch gonadal germ cell clusters represent developmentally distinct states

To further determine how the zGC1 and zGC2 clusters differ from somatic cells, we assessed differentially expressed genes (DEGs) between the transcriptomes of the zGC clusters and the somatic (zSomatic) gonadal cells, with DEGs defined as genes with expression in ≥10% of cells in the target cluster, a log-fold change ≥ 0.5 and an adjusted p-value < 0.05. Both zGC1 and zGC2 shared 1,077 DEGs relative to zSomatic clusters (524-up and 553-down regulated; Figure 3A; Supplemental table 6); these included other general germ cell markers not noted above, such as *TDRD15*, *PIWIL1*, *MAEL* and *SMC1B* (Figure 3B). Another 1,093 DEGs were identified only for zGC1; these included several canonical PGC pluripotency markers, such as *PRDM14* and *KIT* (Figure 3B) (Magnusdóttir et al., 2013; Srihawong et al., 2016). Notably, these canonical PGC markers were largely absent or lowly expressed in zGC2.

**Figure 3.**
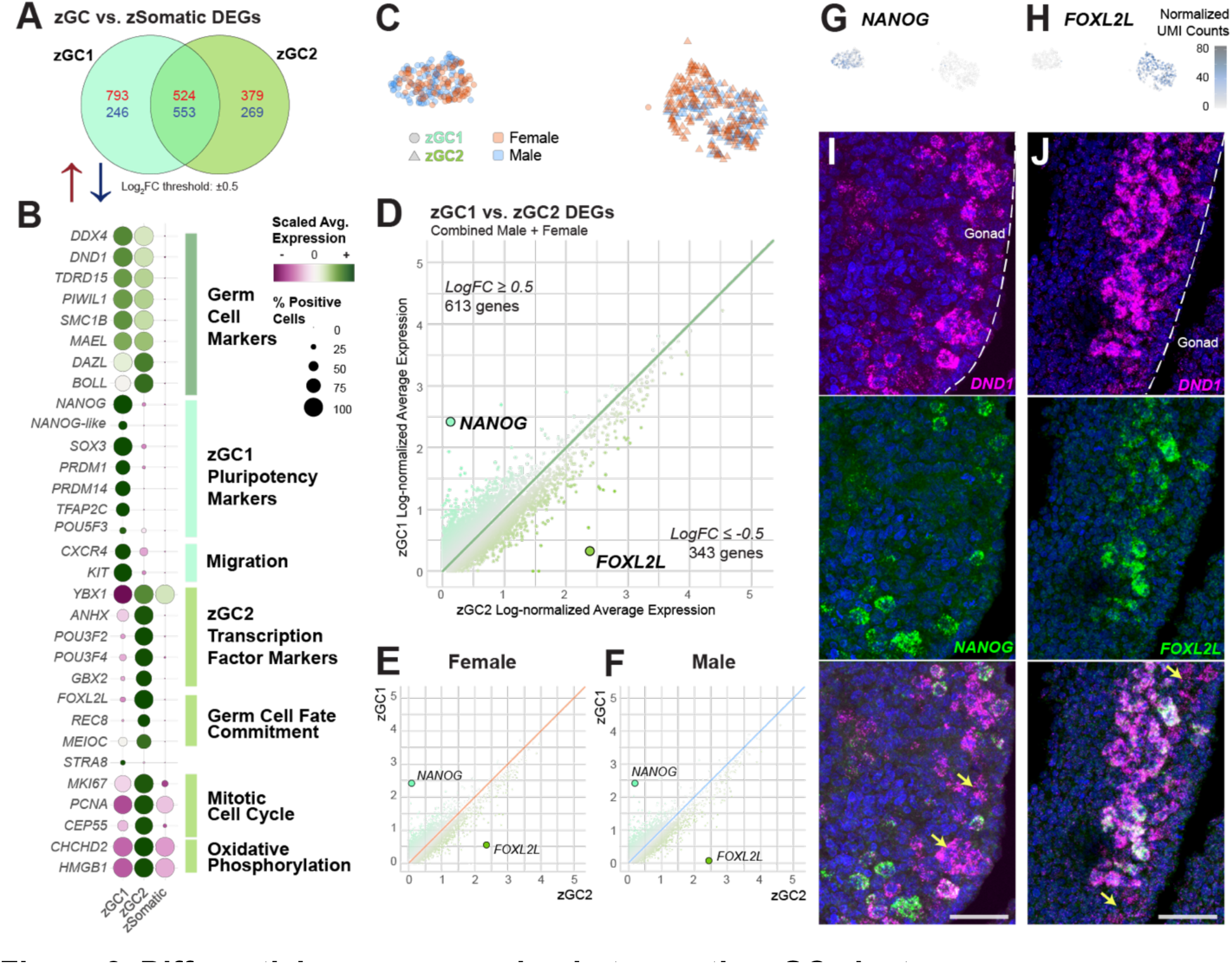
Differential gene expression between the zGC clusters. A. Venn diagram of upregulated (red) and downregulated (blue) gene expression between each zGC cluster and all zebra finch somatic cell types (zSomatic). Differential expression gene (DEG) threshold is defined as a log-fold change cutoff at ±0.5, percent expressing cells > 10%, and an adjusted p-value ≤0.05. B. Dotplot of select gene marker scaled expression between zGC and aggregate zSomatic clusters, with broad gene annotations listed to the right. C. Abridged UMAP plot of zebra finch zGCs, highlighting the corresponding cell barcode sex by color and germ cell type by shape. D. Log-normalized gene expression of zGC1 (y-axis) and zGC2 (x-axis) clusters for each gene. Points are colored by the relative log-fold change in gene expression between clusters, with the most differential genes, *NANOG* (LOC100230680) and *FOXL2L* (LOC101233936), highlighted. E. Log-normalized gene expression of male zGC1 (y-axis) and zGC2 (x-axis) clusters, separated by sex. *NANOG* and *FOXL2L* are highlighted. F. Log-normalized gene expression of female zGC1 (y-axis) and zGC2 (x-axis) clusters. *NANOG* and *FOXL2L* are highlighted. G. zGC UMAP overlaid with *NANOG* expression (transcript UMI/10,000 total cell UMIs) in each cell barcode. H. zGC UMAP overlaid with *FOXL2L* expression (transcript UMI/10,000 total cell UMIs) in each cell barcode. I. Dual-label *in situ* hybridization of germ cell marker *DND1* and *NANOG*. Yellow arrows highlight *DND1*+ cells without *NANOG* signal. Scale bar = 50µm. J. Dual-label *in situ* hybridization of germ cell marker *DND1* and *FOXL2L*. Yellow arrows highlight *DND1*+ cells without *FOXL2L* signal. Scale bar = 50µm.

We identified 648 DEGs between zGC2 and zSomatic clusters; these included several homeobox (e.g., *YBX1, GBX2, DLX2*) and POU domain (e.g., *POU3F2, POU3F4*) transcription factors (Figure 3B). zGC2 also showed strong upregulation of fate determination markers *MEIOC, REC8* (*LOC121468792*), and *FOXL2L* (*LOC101233936*). *FOXL2L* (alternatively *FOXL3-like*) has been identified as a cell-intrinsic suppressor of spermatogenesis in medaka fish (Nishimura et al., 2015) and a driver of oogonial progenitor cell fate determination in zebrafish (Liu et al., 2022) that corresponded with increased cell proliferation. We noted that many zGC2 DEGs also had roles in mitotic cell cycle (*MKI67, CDCA3, PCNA*, *CEP55*) and oxidative phosphorylation pathways (*HMGB1*, *CHCHD2*) (Aras et al., 2015; Tang et al., 2011), both of which occur during cell proliferation (Yao et al., 2019). Indeed, cell cycle scoring indicated that 55% of zGC2 cells were in the G2 or M phase compared to 22% of zGC1 cells (Figure 1–figure supplement 2D; Supplemental table 3). This difference persisted despite cell cycle regression during the clustering workflow.

Looking at only the zGC clusters, we visually confirmed significant representation of zGC clusters in the male and female datasets (Figure 3C). Between zGC1 and zGC2 we identified 956 DEGs, with the most distinct markers for each cell populations being *NANOG* for zGC1 and *FOXL2L* for zGC2 (Figure 3D; Supplemental tables 8 and 9), and this persisted for each sex (Figure 3E-F). These markers appeared mutually exclusive by UMAP (Figures 3G-H). We assessed these markers *in vivo* by fluorescent dual-label *in situ* hybridization on transverse sections of the zebra finch HH28 gonads, finding incomplete co-localization of *NANOG* and *FOXL2L* in *DND1*+ germ cells (Figure 3I-J) and no other cell type. This confirmed these genes as markers of the zebra finch zGC1 and zGC2 cell types, respectively, at HH28. We noted that *NANOG*+ germ cells were generally located toward the posterior and anterior ends of the gonad, while *FOXL2L*+ germ cells were more tightly packed near the center of the medial edge, facing the dorsal mesentery. We additionally noted for both sexes that *FOXL2L* expression was found in *DND1*+ germ cells of both left and right gonads (Figure 3–figure supplement 1).

Taken together, these gene expression marker findings imply that the zGC1 cluster is in a stem cell state, while zGC2 is in a fate determination and proliferative expansion state. Notably, we found that this heterogeneity exists in both sexes (Figure 3C). In the broader context of germ cell developmental stages across vertebrates, we infer zGC1 as being gonadal PGCs and zGC2 as pre-meiotic gonial progenitor cells, respectively falling on earlier or later gametogenic timepoints.

### Sex-biased gene expression in zebra finch gonadal germ cell clusters

While we found that differences between zGC1 and zGC2 were not predominantly due to sex differences (Figure 3C and Figure 1–figure supplement 2C), further analyses revealed some minor sex differences within cell populations (Figure 3–figure supplement 2A). Interestingly, there were twice as many DEGs between male and female zGC1 (n=203) than zGC2 (n=102; Figure 3–figure supplement 2A; Supplemental tables 10 and 11), despite zGC2 expressing more markers of sexual fate determination. Many of these DEGs were sex chromosome genes (zGC1: n=85; zGC2: n=67). Nonetheless, there were fewer DEGs between sexes than those found between the zGC clusters (956 DEGs; Supplemental table 6) and several of the top zGC markers were expressed in both sexes at roughly equal levels (Figure 3–figure supplement 2B).

### Re-analysis of an independent dataset supports two germ cell populations

A previously published study using single-cell datasets of male and female zebra finch embryonic gonads at HH28 identified three “PGC subtypes” that they defined as: 1) high pluripotency; 2) high germness; and 3) low germness/pluripotency. (Jung et al., 2021; denoted Seoul National University (SNU) dataset relative to our Rockefeller University (RU) dataset). We sought an explanation for the differences of the number of clusters and their cell type substates between studies. As their analysis did not incorporate several standard quality controls that we used here, we reprocessed their datasets before and after applying much of our quality control workflow. Indeed, when we incorporated ambient RNA removal and mitochondrial genome mapping, but only removed cells expressing ≤ 200 genes as in their study, we actually inferred four (instead of two or three) germ cell clusters (c13, c17, c22, c29; Figure 1–figure supplement 3A). However, we noted a bimodal distribution in summary statistics of the SNU datasets (Figure 1–figure supplement 3B), and that clusters c13 and c29 had much lower UMI and gene counts than the other two clusters, and c29 additionally had high mitochondrial gene expression (Figure 1–figure supplement 3C). When we applied the appropriate quality control filters (Supplemental table 1), 39.4% of the SNU cell barcodes were removed (compared to 14.9% equivalently removed in our RU dataset; Figure 1–figure supplement 3E vs. 3F), and among the removed barcodes labeled as germ cells, most were derived from the c13 and c29 clusters (Figure 1–figure supplement 3G). A large portion of removed cells were erythrocytes (Figure 1–figure supplement 3E).

After quality control filtering of the SNU dataset, the remaining cells generated a UMAP landscape of gonadal cell types similar to our dataset (Figure 1–figure supplement 4A; Supplemental table 12). Importantly, this analysis left only two germ cell clusters remaining, primarily made up of barcodes from c17 and c22 in the unfiltered dataset (Figure 1–figure supplement 3G); now labelled as c10 and c25 in the filtered dataset (Figure 1–figure supplement 4A). A comparative reference-query mapping and label transfer analysis (Stuart et al., 2019) of the filtered SNU dataset to the filtered RU dataset showed high concordance between expression profiles of the clustered cell types (Figure 1–figure supplement 4B). The c10 and c25 SNU filtered dataset analyses matched the distinct zGC1 and zGC2 clusters of the RU dataset. Importantly, we found similar DEG markers for these clusters (Figure 1–figure supplement 4B; Supplemental table 13), and similar module score enrichments for the candidate GRC gene paralogs in the SNU zGC populations (Figure 1–figure supplement 4D; Supplemental table 7). These findings across independently generated scRNAseq datasets support two distinct but closely related clusters in the zebra finch gonad at HH28.

### Single-cell transcriptomic analysis identifies one germ cell population in the HH28 chicken gonad

To compare zebra finch and chicken, we generated scRNAseq datasets from male (n=2) and female (n=2) chicken embryonic gonads at HH28, a stage where chicken PGCs are commonly collected for assisted reproductive technology applications (Choi et al., 2010). This stage occurs just prior to the HH29 sexual differentiation of developing gonadal tissue (Ayers et al., 2015; Estermann et al., 2020b). The chicken samples were processed simultaneously and with the same quality control steps as the zebra finch samples (Figure 4–figure supplement 1; Supplemental table 1). A total of 8,607 cells were mapped against a chicken reference genome with 24,180 gene annotations (GCF_000002315.6; Supplemental table 14) and visualized by UMAP (Figure 4A; Supplemental table 15). Clustered cell types were identified through nearest-neighbor clustering and marker-based label transfer (Figure 4B). Between chicken datasets we noted a higher total number of female cells than male, but cell type proportions between sexes remained roughly equivalent (Figure 4–figure supplement 1D). These cell types were similar to those found in the zebra finch, as they broadly shared many of the same gene markers (Figure 4C vs Figure 1D; Supplemental table 14).

**Figure 4.**
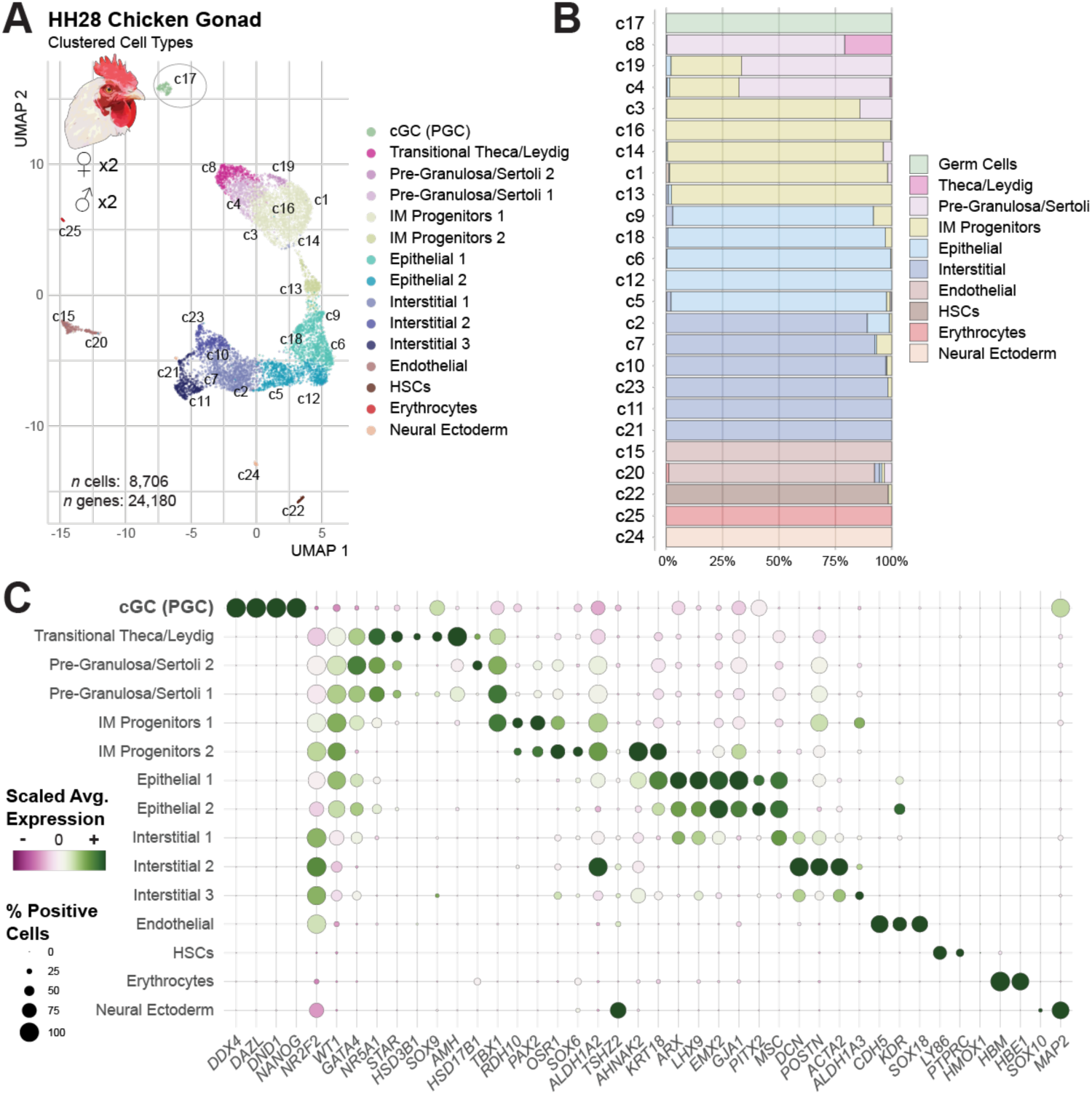
The chicken HH28 gonad demonstrates only one germ cell cluster. A. UMAP plot of male and female chicken gonadal at HH28. Cells are colored by the clustered cell type, with initial nearest-neighbor cluster labels overlaid. Further information on quality control and dimensional reduction for this dataset may be found in Figure 4– figure supplement 1. B. Proportional bar chart of inferred cell types in each nearest-neighbor cluster. C. Dotplot of scaled expression for select gene markers of each clustered cell type identified in the HH28 chicken gonad.

In contrast to the zebra finch, only one chicken germ cell (cGC) cluster was found (c17, Figure 4A-C) and it remained stable across multiple clustering resolutions (Figure 4–figure supplement 1B). An assessment of DEGs between cGCs and chicken somatic (cSomatic) cells marked the cGC cluster with 1,049-up regulated and 380-down regulated genes. The up-regulated genes included many canonical PGC markers, such as *NANOG*, *POU5F3* (*OCT4* homolog), and *KIT* (Figure 4–figure supplement 2A; Supplemental table 16). To validate a unitary PGC population, we demonstrated a complete overlap of *DAZL* and *NANOG* in HH28 chicken gonads and dorsal mesentery by fluorescent *in situ* hybridization (Figure 4–figure supplement 3). Between male and female chicken cGC clusters, there were 2-3 times fewer DEGs than seen for either zGC cluster (n=59; Figure 4–figure supplement 2B; Supplemental table 17), with about half of these genes located on the sex chromosomes (n=27). Consistent with prior studies (Rengaraj et al., 2022), these results support the presence of just one germ cell state in the chicken gonad at HH28, which we identify to be gonadal PGCs.

### Comparison of chicken and zebra finch HH28 gonadal germ cells

To directly compare the chicken and finch HH28 gonadal cells, we integrated the processed RU datasets using 13,913 identified orthologous gene pairs between species (Supplemental tables 2 and 14). A reference-query label transfer analysis of the clustered cell types showed good mapping between the cell types (Figure 5A; Figure 5–figure supplement 1A); though the Mesenchymal “supercluster” (IM Progenitors, Pre-Granulosa/Sertoli and Theca/Leydig cell types) showed lower overlap between species. Of note was a higher proportion of IM progenitor cells versus pre-Sertoli and Granulosa cells in the chicken compared to the zebra finch (Figure 5B), matching previously published findings (Estermann et al., 2021). Other cell types of each species, such as the endothelial and epithelial cell clusters, largely conformed to roughly equivalent general UMAP coordinates (Figure 5B).

**Figure 5.**
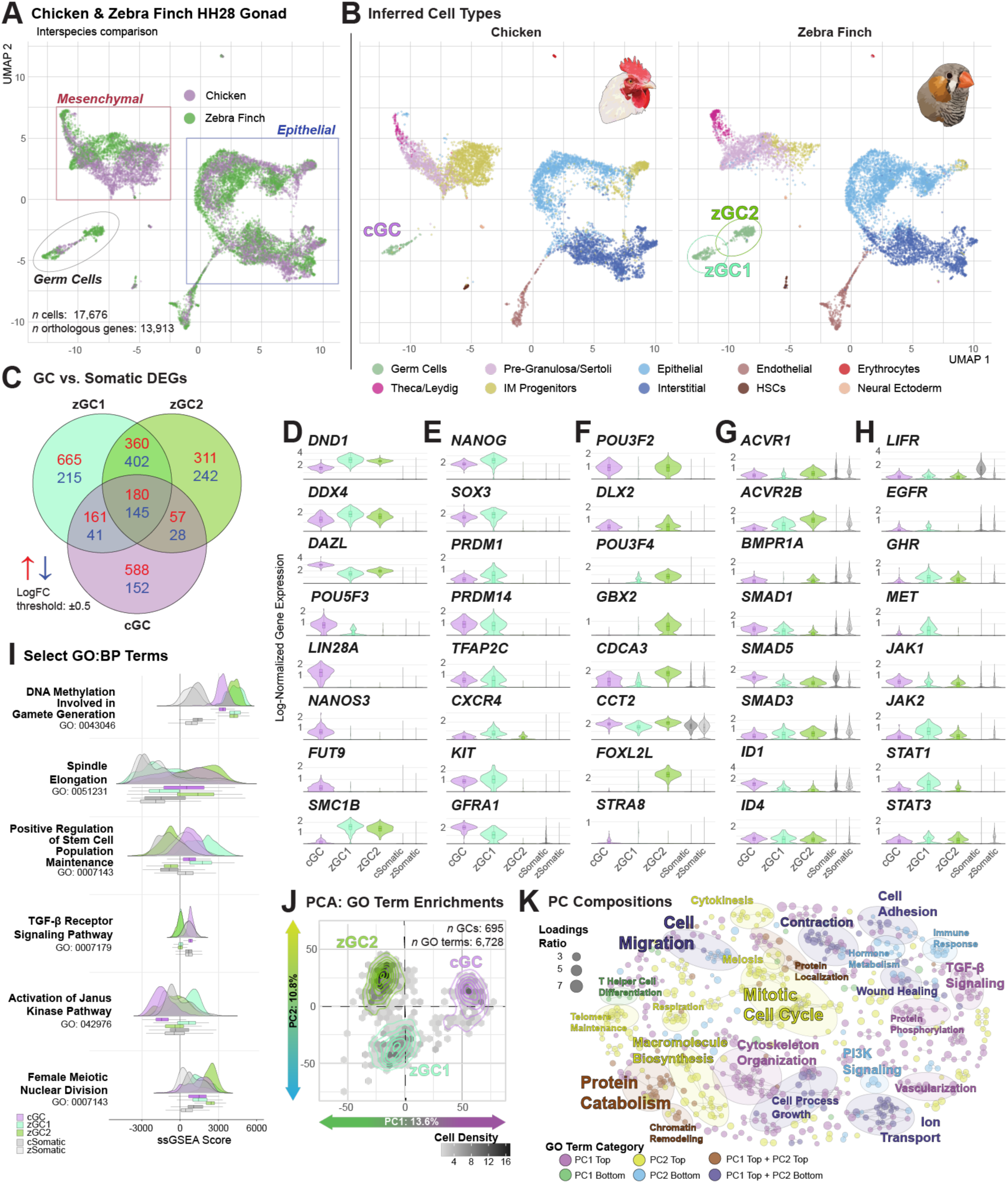
Comparison of chicken and zebra finch HH28 germ cell clusters. A. UMAP plot of integrated chicken (purple) and zebra finch (dark green) gonadal datasets at HH28. B. Separation of the integrated UMAP in subfigure A by species and colored by inferred cell type. C. Venn diagram of upregulated (red) and downregulated (blue) gene expression between each GC cluster and species-respective somatic cell types. A differential expression threshold is defined at a log-fold change of ±0.5. D-H. Violin plots of log-normalized gene expression between GC clusters from each species. Aggregate somatic expression for chicken (cSomatic) and zebra finch (zSomatic) are provided in grey. I. Ridge plots of select GO Terms, showing relative single-sample gene set enrichment analysis (ssGSEA) scores between GC and somatic cell barcodes. J. Projection of Principal Component Analysis (PCA) for all GO Term enrichments assessed for each GC cluster. K. EMAP Plot highlighting principal component GO Term loadings connected by Jaccard score. See supplemental Table 21 for cluster compositions.

A comparison of the germ cell clusters for each species revealed that the chicken cGC clustered with the zGCs rather than with the other somatic cell types (Figure 5A). For the germ cells in the integrated UMAP, the chicken cGC occupied an intermediate position between zGC1 and zGC2 (Figure 5B; Figure 5–figure supplement 1A; Supplemental table 18). Nearest-neighbor clustering of the integrated species dataset identified two germ cell clusters, c20 and c21, with c20 primarily composed of both cGC and zGC1 cells and c21 almost exclusively composed zGC2 cells (Figure 5–figure supplement 1B-D). Examining individual DEGs between germline and species-specific somatic clusters (Figures 5C-H), both zebra finch and the single chicken germ cell populations shared many marker genes (n=325; Figure 5C), including *DND1*, *DDX4,* and *DAZL* (Figure 5D; Supplemental table 19). Consistent with the clustering analyses, the cGC and zGC1 populations shared upregulated gene expression of many pluripotency markers, including *NANOG, SOX3, PRDM1, PRDM14*, and *TFAP2C* (Figure 5E) (Chambers et al., 2007; Jean et al., 2015; Magnúsdóttir et al., 2013; Motono et al., 2008), migratory markers *CXCR4* and *KIT* (Lee et al., 2017; Srihawong et al., 2016), as well as the spermatogonial stem cell marker, *GFRA1* (Buageaw et al., 2005). cGC cells also expressed a few genes upregulated in the zGC2 population, such as *POU3F2* and *DLX2*, and several cell cycle genes, such as *CDCA3 and CCT2* (Figure 5F).

In addition to cell identity markers, we identified several growth factor receptor similarities and differences between the three germ cell populations. In all three populations (cGCs, zGC1, and zGC2), there was consistent upregulation of several SMAD and TGF-b superfamily signaling receptors (*ACVR2B*, *SMAD5 and SMAD3*; Figure 5G), though *ACVR2B* and *SMAD5* were more highly expressed in zGC2 than zGC1 (Supplemental table 6). However, compared to the cGC cluster, zGC2 demonstrated poor expression of *SMAD1*, and receptor subunit genes *ACVR1* and *BMPR1A* were notably downregulated in the zGC1 cluster. These findings suggest that BMP and Activin signaling within the TGF-beta superfamily, necessary for the maintenance and self-renewal of migration-competent chicken PGCs (Whyte et al., 2015), may have divergent roles in zebra finch germ cell development.

We noted some clear species differences. Several well-characterized chicken germ cell markers, *POU5F3, LIN28A, NANOS3*, and *FUT9* (an SSEA-1 epitope synthesis gene) had low or absent expression in both zebra finch zGC1 and zGC2 (Figure 5D). Conversely, *SMC1B*, a previously identified zebra finch germ cell marker (Jung et al., 2021), was found in both zGC clusters, but low in cGC (Figure 5D). In zGC1, we also found significant upregulation of several JAK/STAT-related receptors (e.g., *GHR, MET)* and downstream genes (e.g., *JAK2*, *STAT1*) not upregulated in cGC (Figure 5H). Importantly, only zGC2 expressed fate determination markers, such as *FOXL2L* (Figure 5F), but, interestingly, did not have significant expression of *STRA8* (Figure 5F), an RA-stimulus response gene canonically signaling the onset of meiotic fate determination in chicken (Smith et al., 2008). To ensure that the absence of expression was not due to annotation error, raw read alignments for several orthologs with species-specific expression were manually reviewed against their respective genome references (Figure 5–figure supplement 2). We found no evidence of annotation or other error to explain these species differences.

We wondered whether the cGC cluster shared any expression with the identified GRC gene paralogs, as found in the zGC clusters. We scored gene modules composed only of zebra finch GRC gene candidates with chicken paralogs (n=69) and saw no major enrichment in cGC vs. cSomatic clusters (Log2FC<0.5; Figure 5–figure supplement 3A; Supplemental table 7). The zebra finch module enrichments were similar between the orthologous geneset and the full geneset (Figure 5–figure supplement 3B vs. Figure 2C). In particular, we also found that chicken *NAPA* was not upregulated in cGC vs. cSomatic clusters, like zebra finch *NAPAA* but not *NAPAGRC* (Figure 5–figure supplement 3C). Altogether these results imply that zebra finch GRC genes provide unique germline expression patterns not demonstrated by either the zebra finch A chromosome paralogs or chicken A chromosome orthologs.

### Functional gene category differences between zebra finch and chicken primordial germ cells

To assess broader functional characteristics between the germ cell populations, we ran single-sample gene set enrichment analysis (ssGSEA) against 6,728 Biological Process Gene Ontology terms (GO; Aleksander et al., 2023) containing more than five zebra finch/chicken gene orthologs (Supplemental table 20). As expected, each germ cell cluster was enriched for several germ cell-related GO terms compared to gonadal support cell populations, including “DNA Methylation Involved in Gamete Formation” (Figure 5I). Compared to somatic cell enrichments, mitotic cell division terms (e.g., “Spindle Elongation”) were enhanced in zGC2 and cGC, while terms such as “Positive Regulation of Stem Cell Population Maintenance” were enhanced in zGC1 and cGC (Figure 5I). Interestingly, cGCs but not zGCs were enriched for “TGF-beta Receptor Signaling Pathway” compared to their corresponding somatic cells, whereas zGC1 was exclusively enriched for “Activation of the Janus Kinase Pathway,” mirroring the individual DEG observations. Notably, only zGC2 was enriched for “Female Nuclear Meiotic Division.”

We applied PCA for all GO enrichment scores for the germ cell populations across 694 PCs (Supplemental table 21), with PC1 and PC2 accounting for 13.6% and 10.8% of the variation, respectively (Figure 5J). More than 90% of the total variance was accounted for by PC3-PC375, though none individually accounted for more than 4% of the total variation. PC1 primarily acted to delineate species differences, while PC2 separated the zGC1 and zGC2 populations (Figure 5J).

To identify larger trends between the three germ cell populations, GO terms contributing most to PC1 and PC2 were projected onto an enrichment map, and clustered by Jaccard similarity. The identified PC1 terms had a notable right-sided contribution bias (355 positive terms; 14 negative terms) and had broad enrichment categories differences in TGF-b superfamily signaling, vascularization, and cytoskeletal organization (top quadrant) and T helper cell differentiation (bottom quadrant) (Figure 5K; Supplemental table 22). Terms on the opposing ends of PC2 (231 positive terms; 121 negative terms) resolved clusters broadly defined by mitotic cell cycle (top quadrants) and macromolecule biosynthesis terms and cell migration (bottom quadrants). We also saw cluster differences for GO terms involved in JAK/STAT, PI3K/AKT, and WNT signaling pathways. Overall, these species and germ cell type functional differences support a distinction in all three populations and highlight the complex and dynamic nature of germ cell populations in avian embryonic gonads.

### Cross-species functional analysis of gonadal somatic cells

Considering the developmental differences between chicken and zebra finch germ cell clusters, we sought to assess functional differences of particular extrinsic signaling pathways in the developing gonadal somatic cells. We found species differences in gene expression between markers of sex hormone biosynthesis (Figure 5–figure supplement 4). Namely, the zebra finch mesenchymal cell “supercluster” (Figure 5–figure supplement 4A), and to a lesser extent the epithelial supercluster, showed upregulated expression of sex hormone synthesis genes (Figure 5–figure supplement 4B). Compared to chicken, the *HSD3B1* progesterone biosynthesis enzyme gene was elevated in zebra finch mesenchymal and epithelial clusters. ssGSEA highlighted an enrichment of “Progesterone Biosynthetic Process” (GO: 0006701) in zebra finch somatic clusters compared to chicken (Figure 5–figure supplements 4C). Germ cells of both species expressed the nuclear progesterone receptor (*PGR*) and several membrane progesterone (*PAQR3*, P*AQR8*) receptor genes (Figure 5–figure supplements 4D). The *HSD17B1* redox enzyme gene that enhances androgen and estrogen potency was also elevated in zebra finch clusters, though androgen and estrogen receptors were not highly expressed in any zGC or cGC clusters at this stage. These hormones have critical roles in sex determination of the developing avian gonad (Ayers et al., 2013; Clinton and Zhao, 2023; Smith et al., 2009).

We identified differences for retinoic acid (RA) signaling (GO: 0042573 “Retinoic Acid Metabolic Process”), which was more highly enriched in chicken somatic cells compared to zebra finch (Figure 5–figure supplement 5A). Indeed, compared to zebra finch, chicken somatic cells demonstrated higher gene expression of *ALDH1A2*, whose protein product converts retinaldehyde into RA, and lower levels of the *CYP26B1* retinoic acid degradation gene (Figure 5–figure supplement 5B). Interestingly, while *STRA8* was absent in all germ cell clusters (Figure 5F), both zGC clusters showed higher expression of several RA signaling and stimulus response genes not elevated in the cGC cluster (e.g., *OPN3*, *RBP5, STRA6*, *RARB*; Figure 5–figure supplement 5C).

Taken together, these findings suggest that the somatic cells of the zebra finch gonad begin sexual differentiation of the gonads by HH28, while the chicken gonads remain in a bipotential state, prior to ovarian or testicular commitment starting at HH29 (Ayers et al., 2015; Estermann et al., 2020a, 2020b; Smith et al., 2008). Moreover, the expression patterns of RA biosynthesis and response genes suggest key species differences in the sensitivity and timing of RA signaling in gonadal development between chicken and zebra finch.

### Gonadal *FOXL2L* expression occurs in zebra finch as early as HH25

We sought to further assess zebra finch germ cell heterogeneity *in vivo* across multiple stages of gonadal development through dual-labeling of *NANOG* and *FOXL2L*. In addition to both male and female zebra finch HH28 gonads, each germ cell marker could be distinguished in cells, without co-localization, at earlier (HH25) and later stages (HH36; Figures 6ª-C), documenting germ cell heterogeneity at multiple developmental timepoints. This finding at HH25 was particularly unexpected, as *NANOG+* PGCs were still found in the dorsal mesentery (DM) and potentially migrating toward the gonadal ridge. This was further supported by incomplete co-localization of *DND1*+ germ cells with *NANOG* (Figure 6–figure supplement 1) or *NAPAGRC* (Figure 6–figure supplement 2) at this stage. In sections of HH36 zebra finch gonads, we generally saw many more *FOXL2L*+ cells than *NANOG+* cells, though both populations could be confidently identified in each sex. These data suggest that germ cell fate determination marked by *FOXL2L* readily occurs upon zebra finch germ cell settlement into the gonadal ridge, and that the proportion of these cells increases over development.

**Figure 6.**
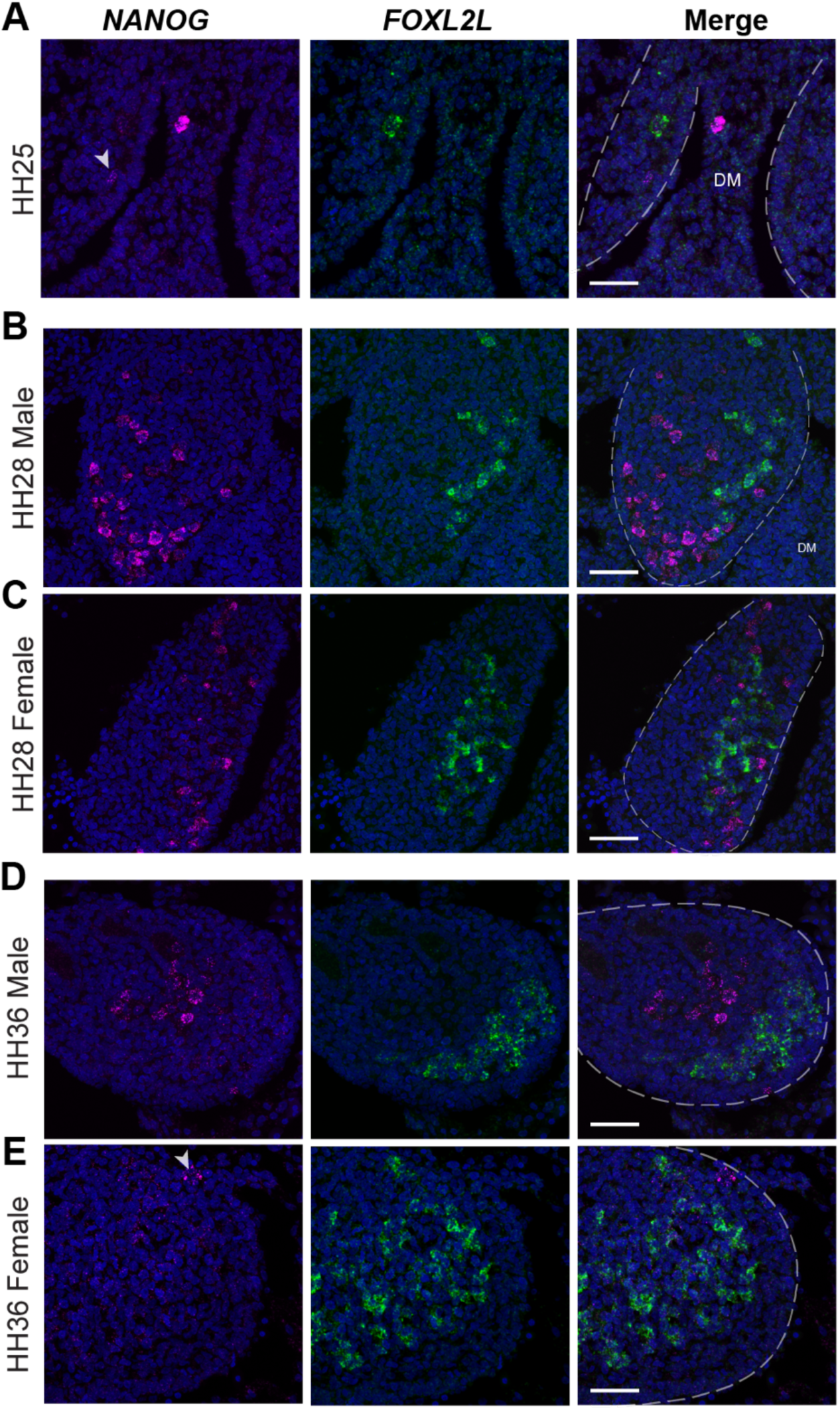
Zebra finch germ cell heterogeneity across gonadal development. A. Dual-label *in situ* hybridization of *NANOG* and *FOXL2L* in HH25 gonads. Arrowhead denotes *NANOG* signal in gonad. B. Dual-label *in situ* hybridization of *NANOG* and *FOXL2L* in HH28 male gonads. C. Dual-label *in situ* hybridization of *NANOG* and *FOXL2L* in HH28 female gonads. D. Dual-label *in situ* hybridization of *NANOG* and *FOXL2L* in HH36 male gonads. E. Dual-label *in situ* hybridization of *NANOG* and *FOXL2L* in HH36 female gonads. Arrowhead denotes *NANOG*+ signal in gonad. F. Scale bars = 50µm. White dotted lines denote gonadal boundary; DM = Dorsal Mesentery.

### Zebra finch germ cell heterogeneity parallels that of HH36 chicken females

To compare gonadal germ cell differentiation between species and potentially identify similar gene expression profiles to zGC2 in chicken, we utilized previously published scRNAseq datasets of chicken embryonic gonadal development where germ cell expression patterns had not been extensively explored (Estermann et al., 2020; denoted as MU for Monash University). We processed the MU datasets using our analysis workflow to assess germ cell development across multiple timepoints: HH25 (their embryonic day 4.5 (E4.5)), HH30 (E6.5), HH35 (E8.5), and HH36 (E10.5; Figure 7–figure supplement 1; Supplemental table 23). To assess the batch comparability of the RU and MU datasets, we compared our RU HH28 chicken datasets to the closest MU time point, male and female HH30. We found that our inferred cell type classifications largely matched the somatic cell type classifications used by the MU study (Figure 7–figure supplement 2A-B). The mesenchymal supercluster showed less distinct similarities, with the HH30 IM progenitor population much smaller proportionally than that found at HH28 (Figure 7–figure supplement 2B). This analysis concurs with the known timing (HH29; Ayers et al., 2015) of sexual differentiation in the chicken gonad.

**Figure 7.**
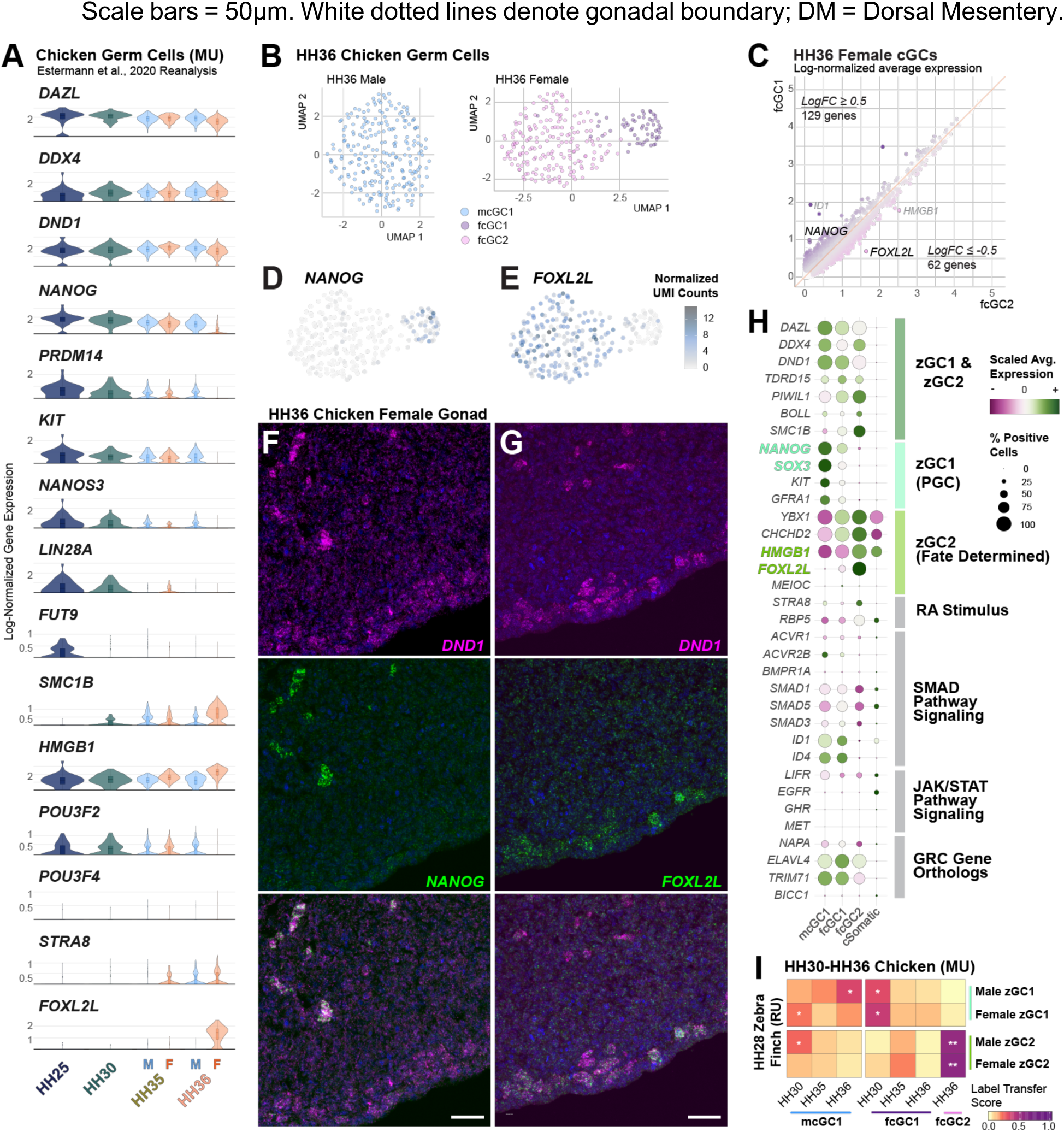
Chicken germ cell heterogeneity in later embryonic development. A. Violin plots of select genes in male and female chicken gonadal germ cells on different embryonic days. B. Individual subclustering of male and female HH36 chicken germ cells. Note the one cluster resolved in the male dataset vs. the two clusters in the female dataset. C. Comparison of average log-normalized gene expression between fcGC1 (y-axis) and fcGC2 (x-axis). A selection of the highest log-fold change genes are labeled. D. HH36 Female cGC UMAP overlaid with *NANOG* expression (transcript UMI/10,000 total cell UMIs) in each cell barcode. E. HH36 Female cGC UMAP overlaid with *FOXL2L* expression (transcript UMI/10,000 total cell UMIs) in each cell barcode. F. Dual-label *in situ* hybridization of germ cell marker *DND1* and *NANOG* in chicken female HH36 gonads. Scale bar = 50µm. G. Dual-label *in situ* hybridization of germ cell marker *DND1* and *FOXL2L* in chicken female HH36 gonads. Scale bar = 50µm. H. Dotplot of select gene marker scaled expression between E10.5 male and female cGCs and aggregate cSomatic clusters. Gene symbols highlighted by color correspond to zGC1 (teal) or zGC2 (lime) marker conservation. I. Confusion matrix of label transfer similarity scores for male and female zebra finch zGC clusters (RU) against chicken (MU) germ cells at HH30, HH35, and HH36. A log2FC>0.50 against other MU stage scores and a p-value≤0.05 by one-sided t-test is denoted by *. A log2FC>2.0 is denoted by **.

Notably, in an aggregate of all MU datasets as well as for each male and female chicken gonadal time point, our analyses resolved only one cluster of germ cells (Figure 7–figure supplement 1B-C). cGCs showed progressive declines in gene expression of several stem cell markers (e.g., *NANOG, PRDM14*, *LIN28A)*, though sizeable expression only persisted in the male HH36 gonadal dataset (Figure 7A). In contrast, several genes showed differential expression patterns between female HH35 and HH36 germ cells (Figure 7A; Supplemental table 24), corresponding with the RA-mediated onset of oogenesis in chicken around this developmental stage (Rengaraj and Han, 2022; Smith et al., 2008). The loss of *NANOG* and other pluripotent markers coincided with *FOXL2L* expression in female HH36 germ cells, matching the known onset of *FOXL2L* upregulation in the left gonad of female chicken embryos at E9 (Ichikawa et al., 2019).

As only one cluster of chicken germ cells was derived at each stage, we sought to discern any germ cell heterogeneity within the HH36 scRNAseq datasets. By individually subclustering the HH36 chicken germ cells for each sex, we resolved two female germ cell clusters that we denoted as fcGC1 and fcGC2 (Figure 7B; f for female). In contrast, the male cells still formed only one *NANOG*+ cluster (mcGC1; Figure 7B; Supplemental table 25). The female clusters were distinguished from cSomatic clusters and each other by several markers, notably *NANOG* (fcGC1) and *FOXL2L* (fcGC2) (Figure 7C-E; Figure 7–figure supplement 3A; Supplemental tables 25-27). Dual-label *in situ* hybridization validated these patterns in chicken HH36 gonads, showing regional exclusivity of *NANOG* and *FOXL2L* gene expression in *DND1*+ cells (Figure 7F-G; Figure 7–figure supplement 4A). *FOXL2L* was not expressed in male HH36 gonads (Figure 7–figure supplement 4B), nor at earlier chicken gonadal stages (Figure 7–figure supplement 4B-D). Between male and female cGC1 clusters at HH36, there were relatively few other genes demonstrating high log-fold change differences, with much of the differential expression coming from sex-chromosome genes (Figure 7–figure supplement 3B; Supplemental table 28).

Between the female fcGC1 and fcGC2 clusters, several differential markers mirrored those found between the zGC1 and zGC2 clusters (Figure 7H; Supplemental table 29). In particular, transcription factors *NANOG* and *SOX3* were highly conserved markers for the zebra finch and chicken female GC1 cluster, while *FOXL2L* and *HMGB1* were consistently upregulated in the female GC2 cluster of both species. Between fcGC1 and fcGC2, several TGF-b/SMAD superfamily signaling pathway genes declined, including those upregulated between zGC1 and zGC2 such as *ACVR2B* and *SMAD5* (Figure 7H). As in the HH28 cGC cluster, JAK/STAT signaling pathway genes were lowly expressed or absent in both fcGC clusters at this stage (Figure 7H). Orthologous GRC gene candidates were also expressed at low levels (Figure 7H), and orthologous GRC module scores also did not demonstrate significant enrichment in the MU cGC clusters (Figure 7–figure supplement 5; Supplemental table 7).

To comprehensively assess corresponding similarities between the chicken and finch germ cell types, we compared the gene expression profiles of all orthologous genes between HH28 zGCs and HH30-36 cGCs by reference mapping analysis. Similarities scores for each zGC-cGC grouping showed male and female zGC1 were diffusely similar to multiple male and female cGC timepoints from HH30 and HH35, but generally paired most closely with cGC populations of their respective sex (Figure 7I; Supplemental tables 30 and 31). In contrast, both the male and female zGC2 populations mapped most closely to female HH36 cGC2 cells. Similar results were found for the zGC clusters in the SNU dataset (Figure 7–figure supplement 6A; Supplemental tables 31 and 32). As a control, an equivalent analysis using the RU chicken datasets mapped the HH28 cGC cells across either MU cGC1 cluster favoring the corresponding sex (Figure 7–figure supplement 6B; Supplemental tables 31 and 33). Collectively, these data show that although chicken PGCs form a relatively uniform population during embryonic development, by HH36 female chicken germ cells begin to segregate into two populations that have similarities to the two finch populations found throughout development.

## Discussion

The study of avian germ cell biology and reproductive development has overwhelmingly focused on chicken and other poultry species, despite the incredible diversity of birds (Flores-Santin and Burggren, 2021; Jarvis et al., 2014). Using scRNAseq datasets in tandem with spatial *in situ* hybridization patterns, we uncovered key differences in the gene expression, sexual dimorphism, and developmental timing of gonadal germ cells between chicken and zebra finch. In particular, two germ cell types exist simultaneously in the zebra finch HH28 embryonic gonad, one we infer as more advanced along the path of germ cell differentiation than the other, while the chicken gonad at the same stage retains a population of one germ cell type. Later in development (HH36), the female chicken gonad demonstrates similarly heterogeneous germ cell populations to the finch, in line with previous observations (Ichikawa et al., 2019; Rengaraj et al., 2022; Smith et al., 2008). Our findings have a host of implications for understanding the evolution of developmental reproductive biology in birds.

The HH28 zebra finch zGC1 and chicken cGC clusters were the most similar to each other, expressing conserved PGC markers of migratory-competence and pluripotency, such as *NANOG*, *PRDM14, CXCR4* and *KIT* (Magnúsdóttir et al., 2013; Okuzaki et al., 2019; Sánchez-Sánchez et al., 2010). However, HH28 zGC1 and cGC also had some fundamental differences, including low expression in zGC1 of *POU5F3* and *LIN28A*, which have critical roles in chicken PGC migration and pluripotency (Meng et al., 2022; Suzuki et al., 2023). These genes precipitously decreased at later developmental timepoints in male and female chicken gonadal germ cells, corresponding to the transition of oocyte formation.

The HH28 zebra finch zGC2 cluster was more dissimilar to the chicken cGC. The notable upregulation of *FOXL2L,* alongside other meiotic onset (*REC8*, *MEIOC*) and proliferative (*PCNA, MKI67*, *HMGB1*) genes with the downregulation of *NANOG*, suggest that the zGC2 population is differentiated from a migratory stem cell state, likely toward pre-meiotic fate determination. The similar *FOXL2L+* germ cell cluster in the later HH36 female gonad coinciding with early oogenesis (Ayers et al., 2015; Smith et al., 2008), corroborates previous reports of *FOXL2L* expression onset in the chicken female left ovary around E9 and peaking at E14 (Ichikawa et al., 2019). This gene is lost in non-placental mammals (Bertho et al., 2016), but has a conserved role in other vertebrates; *FOXL2L* has been identified as a cell-intrinsic suppressor of spermatogenesis in male and female embryos of the medaka fish (Nishimura et al., 2015) and marks the earliest points of germline stem cell commitment to pre-meiotic oocyte progenitors in zebrafish (Liu et al., 2022).

One intrinsic source potentially driving the dramatic differences between chicken and zebra finch germ cell development is the zebra finch GRC. The programmed elimination of the zebra finch GRC during somatic specification and spermatogenesis suggests a unique role for its gene paralogs, potentially to avoid gene regulation conflicts in somatic tissues (Vontzou et al., 2023). Our study appended available gene annotations from a partially sequenced GRC (Biederman et al., 2018; Kinsella et al., 2019), finding significant GRC gene expression differences between zGC1 and zGC2 clusters that did not mirror the expression profiles of their A chromosome counterparts. This germ cell upregulation was also not mirrored by chicken A chromosome orthologs, suggesting that these GRC gene sequences are uniquely regulated in the songbird to provide novel germ cell functions. Future work to fully characterize the GRC and GRC gene roles through sequencing and functional studies will be critical to identify impacts it may have on development of songbird germ cells.

Across vertebrates, germ cell development and differentiation are largely dependent on extrinsic stimuli from the gonadal environment. In the zebra finch HH28 gonad, markers of sex hormone biosynthesis (e.g., *HSD3B1*, *HSD17B1*, *CYP19A1*) were more highly expressed than in the chicken at HH28, consistent with earlier gonadal maturation and sex determination necessary for meiotic onset. This finding aligns with previous work comparing the rate of decline in *PAX2+* IM progenitors in favor of Pre-Sertoli/Granulosa cells, denoting an accelerated maturation of somatic cells in the zebra finch gonad compared to chicken (Estermann et al., 2021). Further characterization of multiple time points of migrating blood and establishing gonadal germ cells, and the extrinsic gonadal cell environment, across avian species, in both sexes, will likely yield even greater diversity of PGC and gonadal states than what we have discovered here.

Interestingly, we did not observe upregulation of *STRA8* in the zGC2 population, which in the HH36 fcGC2 population marks the RA-mediated onset of oogenesis (Bowles et al., 2006; Koubova et al., 2006; Smith et al., 2008). Instead, RA receptors and other markers of RA signaling (e.g., *RBP5*) were expressed in both zGC1 and zGC2 clusters, suggesting another difference in zebra finch and chicken germ cell developmental strategies. Germ cells in several teleost fish species, including the zebrafish and medaka, undergo differentiation independent of *STRA8*, utilizing other signals in tandem with other RA-interacting proteins, such as Rec8a (Adolfi et al., 2021; Crespo et al., 2019). Future work will be necessary to determine if later meiotic stages also occur on an *STRA8*-independent basis in the zebra finch, and whether mechanisms such as those employed in teleosts also exist in the zebra finch.

Beyond developmental biology, our study has important implications for the long-term maintenance of zebra finch PGCs *in vitro*. In chicken, HH28 gonads are used as a source for PGCs for stable cultures (Choi et al., 2010; Han et al., 2002; Shiue et al., 2009; Szczerba et al., 2020), and in previous work we successfully cultured zebra finch gonadal PGCs for several days, injected them in host embryonic gonads, and identified some host gonad colonization (Jung et al., 2019). This highlights the value of embryonic songbird gonads for gene manipulation and biobanking applications. However, these methods produce low yields of migratory-competent zebra finch PGCs and have not enabled long-term cultures. One reason for this could be due to the heterogeneity of gonadal germ cell states we found here, some having already progressed beyond a PGC state. For instance, we identified differential expression of growth factor receptor genes between chicken and zebra finch germ cell clusters, including those in the TGF-beta superfamily signaling pathway, suggesting those factors essential for chicken PGC cultures may not have a conserved role in zebra finch (Whyte et al., 2015). Our findings also predict that zebra finch PGCs may also be more sensitive to progesterone and RA, commonly found in serum and serum replacements. The zGC1 cluster also showed unique upregulation of many genes involved in JAK/STAT signaling. This pathway maintains important roles across many vertebrate stem cell lines, including in chicken spermatogonial stem cells (Herrera and Bach, 2019; Zhang et al., 2015). Recently, short-term cultures of blood-derived zebra finch PGCs have been reported (Gessara et al., 2021), adapting culture conditions used for chicken blood PGCs. As blood-derived PGCs likely represent a purer population with strong migratory cues compared to gonadal PGCs, blood PGCs may be more appropriate for derivation of long-term songbird PGC cultures for germline transmission. Growth factor and small molecule screens of signaling pathway differences between blood and gonadal PGCs could inform the development of long-term zebra finch germline stem cell cultures.

Our studies validated some findings of Jung et al., 2021, on heterogeneity of zebra finch PGCs, as well as differences between chicken and zebra finch (Jung et al., 2023). These include the expression of *SMC1B* in zebra finch but not chicken germ cells, and of stem cell marker expression differences between zGC clusters. However, we find that one of the PGC subtypes, which the authors suggest are cells undergoing biological pruning, is more likely a technical artifact resulting from failure to remove damaged, low quality cells with high mitochondrial DNA content (Osorio and Cai, 2020) and low sequence depths. This is a critical issue in single cell analyses, as not including appropriate UMI and gene count cutoffs can lead to sample artifacts and false discovery in scRNAseq datasets (Ilicic et al., 2016; Luecken and Theis, 2019; Lun, 2018). With proper barcode removal from their dataset, we resolved only two clusters (zGC1 and zGC2), matching what was found in our dataset.

Jung et al. (2023) highlight a potentially enhanced role for Activin signaling in zebra finch PGCs compared to chicken. Consistent with this hypothesis, our analyses show elevated expression of Activin receptors *ACVR1* and *ACVR2B* in the zGC2 cluster compared to zGC1. As this pathway has many dynamic roles across germ cell development (Wijayarathna and Kretser, 2016), we instead predict that cell culture additives supporting Activin signaling in zebra finch PGCs may cause undesirable differentiation and loss of migratory competence.

Our analyses additionally benefitted from the curation of 3’ UTR annotations in the chicken and zebra finch reference genomes. Several of the most utilized scRNAseq library preparations rely on 3’-biased sequencing of mRNA, necessitating adequate gene annotation of those regions to correctly identify expression levels. For instance, our detection of *FOXL2L* gene expression in the zebra finch was the result of our manual curation and extension of NCBI gene annotations, as the default annotation for the zebra finch gene was incomplete (Ichikawa et al., 2019). As more species are studied using single-cell analyses, particularly non-model organisms, the utmost importance must be given to the generation of high-quality reference genomes, such as by the Vertebrate Genomes Project (Rhie et al., 2021), as well as methods to mitigate technical artifacts in cross-species comparisons.

In closing, our study identifies a divergent germ cell developmental program in a songbird, suggesting a far richer diversity in avian germ cell biology than previously identified. One remaining concern is whether the zebra finch GRC or the intensive domestication focus on egg-laying in chicken (Larson and Fuller, 2014; Rubin et al., 2010) facilitated evolutionarily unique quirks of germ cell biology in one or both of these species. Accordingly, the exploration of other representatives across the avian phylogeny will be valuable to determine whether these mechanisms fall along a continuum or represent outliers within the larger clade. This would provide much needed insight on avian germ cell biology necessary for the development of methods for genetic rescue in declining and endangered populations, represented by more than 14% of bird species (IUCN, 2019).

## Acknowledgements

This work was done in part with assistance and equipment from Rockefeller University’s Bioinformatics and Bio-Imaging Resource Centers (RRID:SCR_017791), as well as the Vertebrate Genome Laboratory. Funding for this study was provided by the Howard Hughes Medical Institute, National Science Foundation (EDGE Grant #1645199), and the Revive & Restore Biotechnology for Bird Conservation Program. The authors would like to thank Alexander Suh, John Bracht, Gist Croft, Bruce Draper, Florence Marlow, Carlos Lois, Blanche Capel, Samara Brown, Graham Kelly, Owen Farchione, and Ben Novak for insightful feedback on the data and manuscript.

## Data Availability

Reference genome annotation data will be submitted to public SRA and NCBI databases. scRNAseq datasets will be submitted to GEO, and code for Seurat processing and figure generation will be deposited on GitHub (http://github.com/Neurogenetics-Jarvis and https://github.com/RockefellerUniversity). Accession numbers for public datasets will be provided upon publication. Requests for datasets generated should be directed toward the corresponding authors.

## Author Contributions

MTB, EDJ, and ALK conceived the project. MTB, CS, PC, BH, OF, and ALK generated the sequence libraries. GLG, WW, J-DL, OF, HUT, TC and EDJ provided equipment, reagents, and expertise. MTB, KB, CS, AVS, and ALK performed wet lab experiments. MTB, WW, CS, J-DL, GLG, and TC performed computational analyses. MTB, KB, CS, WW, J-DL, AVS, EH, CR-L, KA- B, TC, EDJ, and ALK analyzed the results. MTB, KB, WW, J-DL, TC, and ALK generated the figures. MTB, EDJ, and ALK wrote the manuscript.

## Methods

### Animal husbandry and sources

Animals were cared for in accordance with the standards set by the American Association of Laboratory Animal Care and Rockefeller University’s Animal Use and Care Committee. Zebra finches were maintained under a 12:12-h light/dark cycle at 18-27°C and breeding pairs provided with a finch seed blend, millet spray, egg mash with fresh squeezed oranges daily, and fresh fruits and vegetables once to twice weekly. A hanging nest box and *ad libitum* jute/cotton mix for nesting material were placed in each cage. Eggs were collected daily and stored at 16-18°C, 80% humidity for up to 7 days. Fertile White Leghorn chicken eggs were obtained from Charles River Laboratories.

### Embryo sexing

Chicken and zebra finch eggs were incubated at 37°C, 60-70% humidity; zebra finch eggs were additionally incubated with intermittent rocking (Showa Furanki). On day 5 of incubation, a small window (2-3 mm diameter) was made in the eggshell of zebra finch eggs, which usually produced a small bleed. Blood was absorbed using Whatman filter paper or a glass needle and then placed into Chelex 100 (Bio-Rad). Chicken eggs were windowed (1 cm diameter) and 1-2 µL blood collected using a glass needle inserted into a vitelline vein. Eggs were resealed with Scotch tape (chicken) or paraffin (zebra finch) and returned to the incubator. DNA was isolated from the blood samples using manufacturer’s instructions for Chelex 100 and sextyping was performed by amplifying the *CHD* genes (primers P2: TCTGCATCGCTAAATCCTTT; P8: CTCCCAAGGATGAGRAAYTG) with a previously published Taq-polymerase PCR protocol (Griffiths et al., 1998).

### Single-cell Collection

On day 6, HH28 embryos were removed from eggs. Gonad pairs were dissected from the embryos using fine forceps and then placed in room temperature 0.05% trypsin-EDTA. Whole gonads were incubated in trypsin for 5 minutes (zebra finch) or 15 minutes (chicken) at 37°C and then dissociated by gently pipetting up and down with a p200 pipette until cell clumps were no longer visible. Trypsin was inactivated with an equal volume of PGC cell culture media containing 10% FBS (Jung et al 2019). For the *in vivo* gonad samples, gonads from two embryos were pooled to create each sample, and four total samples were collected: chicken male and female, and zebra finch male and female. The resulting cells were washed with PGC media and run through a 40µm filter to remove any remaining cell clumps. Samples were resuspended in PGC media and counted using the ThermoFisher Countess II Automated Cell Counter (AMQAX1000) with DAPI vital staining. The following cell counts were obtained for each pooled sample: chicken female ∼700 cells/µl, chicken male ∼1600 cells/µl, zebra finch female ∼2000 cells/µl, zebra finch male ∼2100 cells/µl. Greater than 96% of the cells in each sample were alive.

### Single cell capture on 10x Genomics Chromium

A single Chromium microfluidic Chip B (10x Genomics #2000060) was prepared by pipetting 50% glycerol into all unused wells. A Reverse Transcriptase Master Mix was prepared following the manufacturer’s protocol (Chromium Single Cell 3’ Reagent Kit v3) and was split into four aliquots. Appropriate volumes of water and cell suspension were added to the Master Mix to capture an estimated 7,000 cells for each sample. 10x Genomics v3 GEM Beads (#2000059) and Partitioning Oil (#220088) were then pipetted into the microfluidic chip following manufacturer’s protocol, and a droplet emulsion was created on the chromium instrument. The emulsion was incubated at 53°C for 45 min to allow for reverse transcription and heat deactivated at 85°C 5 min. Emulsion was then broken and cDNA amplified according to manufacturer’s protocol, and the resulting cDNA was measured on a Qubit Fluorometer (ThermoFisher #Q33238). cDNA quantification was as follows: chicken female 12.1 ng/µl, chicken male 17.62 ng/µl, zebra finch female 34.6 ng/µl, zebra finch male 29.6 ng/µl. The resulting cDNA was also visualized on the Agilent Fragment Analyzer (#M5310AA) using the High Sensitivity NGS Kit (#DNF-474-0500) to confirm cDNA size range and primer-dimer prevalence.

### Illumina Library Preparation and Sequencing

cDNA samples were diluted to either 50 ng (chicken) or 100 ng (zebra finch) and were used as input into library preparation for Illumina sequencing following the 10x Genomics protocol (Chromium Single Cell 3’ Reagent Kit v3). Illumina libraries were quantified using a Qubit Fluorometer (#Q33238) and visualized using an Agilent Fragment Analyzer (#M5310AA, DNF- 474-0500). The following quantifications were obtained for the samples: chicken female 22ng/ul; chicken male 27.2 ng/ul; zebra finch female 34 ng/ul; zebra finch male 32 ng/ul. Libraries were labeled using the Chromium i7 Multiplex Kit (PN-120262) and sequenced on either an Illumina HiSeq 4000 or NovaSeq S4 (pair-ended with read lengths of 150 nt) for approximately 2 billion reads per sample.

### Reference genome curation dataset processing

The reference genome and annotation files were downloaded from NCBI (zebra finch: GCF_003957565.2; chicken: GCF_000002315.6). Using previously generated bulk RNAseq datasets, the UTR regions were predicted and added to the annotation file by invoking StringTie (ver 2.1.7). Reference files were built by Cellranger (ver 6.0.1, 10X genomics) mkref command with the polished annotation file, reads were aligned and counted by cellranger count command. Ambient RNA ratios were estimated and cleaned by R package SoupX (Young and Behjati, 2020). Orthologous gene pairs between zebra finch and chicken were identified using BioMart, eggNOG (Huerta-Cepas et al., 2018), reciprocal tBLASTx, and identical gene symbols. All the orthologous genes are listed in (Supplemental tables 3 and 12).

During post-processing analysis, the single-exon zebra finch gene LOC101233936 (*FOXL2L*) was found to be insufficiently annotated, likely due to a high GC-rich region in the 3’ half of the open reading frame (ORF). The annotation was extended through the ORF where a StringTie-identified 3’ UTR was present. Zebra finch datasets were then re-ran against the corrected reference genome and amended LOC101233936 read counts were then added into the existing Seurat objects.

### GRC gene alignment simulation

A simulated small genome was generated as a reference based on the sequence of 8 genes, in which the sequences of 4 genes (*BICC1*, *ELAVL4*, *NAPA*, *TRIM71*) were extracted from the autosomes according to the location of whole genes (UTRs, exons and introns), and the sequences of 4 genes (*BICC1*, *ELAVL4*, *NAPA*, *TRIM71*) from the mRNA sequence of the GRC. Each sequence of these 8 genes were assigned as a chromosome, and a gtf file was generated accordingly. In total, 100,000 96 bp long reads were simulated using the R package Subread based on the reference and gtf files outlined above, which produced a theoretical coverage of 298X (Liao et al 2019). The number of reads per gene was proportional to gene length. Cellranger was used to align the reads back to the genome, and the exon base coverage was calculated by samtools depth (ver 1.12).

### Single-cell RNAseq object processing by Seurat

After ambient RNA removal, the clean matrices were loaded into the Seurat R package (version 4.3.0.1) for downstream analysis. Barcodes falling outside of selected thresholds for ambient RNA-adjusted “nCount_RNA,” “nFeature_RNA,” and the percent mitochondrial genes were removed (Supplemental table 1). Doublet droplets were predicted and removed using doubletFinder (ver 2.0.3).

Seurat objects were normalized and scaled (n=3000 genes) by SCTransform (version 0.3.5; Hafemeister and Satija, 2019) with cell cycle and mitochondrial gene regression. Sample integration, dimensional reduction (n=50 PCs), and nearest-neighbor cell clustering were performed using suggested parameters by the Seurat package. In the RU zebra finch HH28 data, subclustering of zGC2 further resolved clusters corresponding to erythrocytes (c11.8, n=15) and a small number of cells (c11.9, n=12) expressing both hematopoietic stem cell and germ cell markers (Figure 1–figure supplement 2E); c11.9 was excluded from analyses in this study as a potential doublet artifact or an extremely rare population not found by histology (not shown).

Inferred cell types were identified by first identifying reference cells that strictly expressed canonical cell type markers (Supplemental table 2), then using Seurat’s “TransferData” function to identify the cell type identities of the remaining cells based on nearest-neighbor similarity. Clustered cell types were determined by the majority inferred cell type within nearest-neighbor clusters and similar clusters were aggregated using Seurat’s “BuildClusterTree” function. Data visualizations were performed using Seurat functions and modified using ggplot2 commands prior to figure generation in Adobe Illustrator (version 27.5).

Differentially expressed genes were called by the Seurat functions “FindMarkers.” Similarity matrices were generated using the Seurat function “DataTransfer” and the R package ComplexHeatMap (ver 2.12.1). GRC gene module scores were performed using UCell (version 2.0.1; Andreatta and Carmona, 2021). Label transfer and module score significance testing was applied using Welch two sample t-tests, and effect sizes were calculated by log2 fold-change. ssGSEA analysis was performed using the escape R package (version 1.6.0; Borcherding et al., 2021), which utilizes the Molecular Signatures Database 3.0 (Liberzon et al., 2011).

### In situ hybridization

Dual-label *in situ* hybridization was performed using previously published protocols on formaldehyde-fixed embryos. To amplify the genes of interest to be used as probes, briefly, RNA was extracted from zebra finch and chicken embryos using QIAgen RNeasy kit and transcribed into cDNA using LunaScript RT (NEB #E3010). PCR was performed using chicken or zebra finch cDNA and gene specific primers (Supplemental table 33), and Q5 hot start polymerase. PCR products were subsequently cloned into vectors using pGEM-T Easy Vector System II (Promega, Cat# A1380) according to manufacturer’s instructions. Reverse primers were designed with a T3 polymerase binding site for anti-sense transcription. RNA probes were transcribed and labeled with either FITC (fluorescein isothiocyanate) or DIG (digoxigenin) NTPs (Roche Cat#).

Zebra finch and chicken embryos from stages HH25, HH28, HH36 were collected and fixed using 4% PFA and embedded in OCT. Embryos were sectioned using Leica CM 1950 Cryostat at 11µm thickness and preserved on Fisherbrand Superfrost Plus Microscope slides. Dual-label fluorescent *in situ* hybridization (FISH) utilized species-specific probes according to a previous publication’s protocol (Biegler et al., 2021). Slides were counterstained using 1x DAPI, imaged using a Zeiss LSM 780 confocal microscope, and processed using ImageJ (ver 2.0.0-rc-69/1.52p) and Adobe Photoshop CC (ver 24.6.0).

---

**Figure supplement 1.1.**
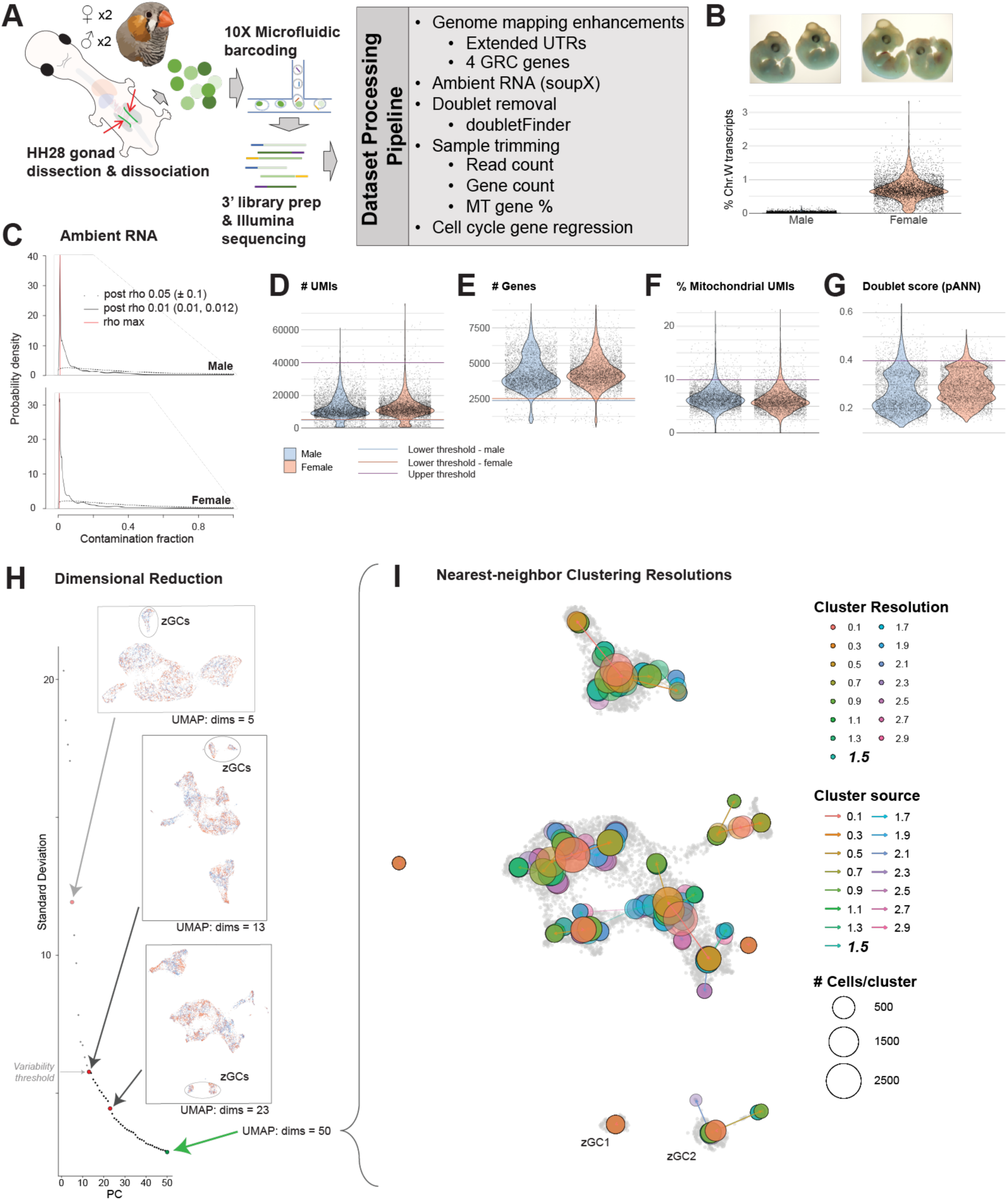
Generation of HH28 zebra finch gonadal datasets. A. Schematic of scRNAseq analysis pipeline, with quality control measures listed. B. Images (top) of HH28 zebra finch embryos used to generate male (left) and female (right) gonadal datasets. Violin plots (below) validate PCR-based sextyping (not shown) by percentage of W sex chromosome genes expressed. C. Graphs demonstrating ambient RNA contamination probability using soupX. Red line denotes rho max normalization of cells. D. Violin plot of UMI counts for all barcodes in male and female datasets, with upper and lower thresholds defined by specified colored lines. E. Violin plot of gene counts of thresholded barcodes (D) for male and female datasets, with lower thresholds defined for each dataset. F. Violin plot of mitochondrial gene percentage of thresholded barcodes (E) for male and female datasets, with at 10% upper threshold cutoff designated. G. Violin plot of doubletFinder artificial nearest-neighbor (pANN) score of thresholded barcodes (F) for male and female datasets, with doublets identified above a score of 0.4. H. Dimensional reduction of trimmed datasets with select UMAP graphs shown for different principal components (PCs). The variability threshold was defined as a change in cumulative variance less than 0.1% between each additional PC. A UMAP representing the first 50 PCs was selected for subsequent analysis. I. Nearest-neighbor clusters at varying resolution, overlaid onto a UMAP representation of the first 50 PCs. A resolution of 1.5 was selected for subsequent analysis.

**Figure supplement 1.2.**
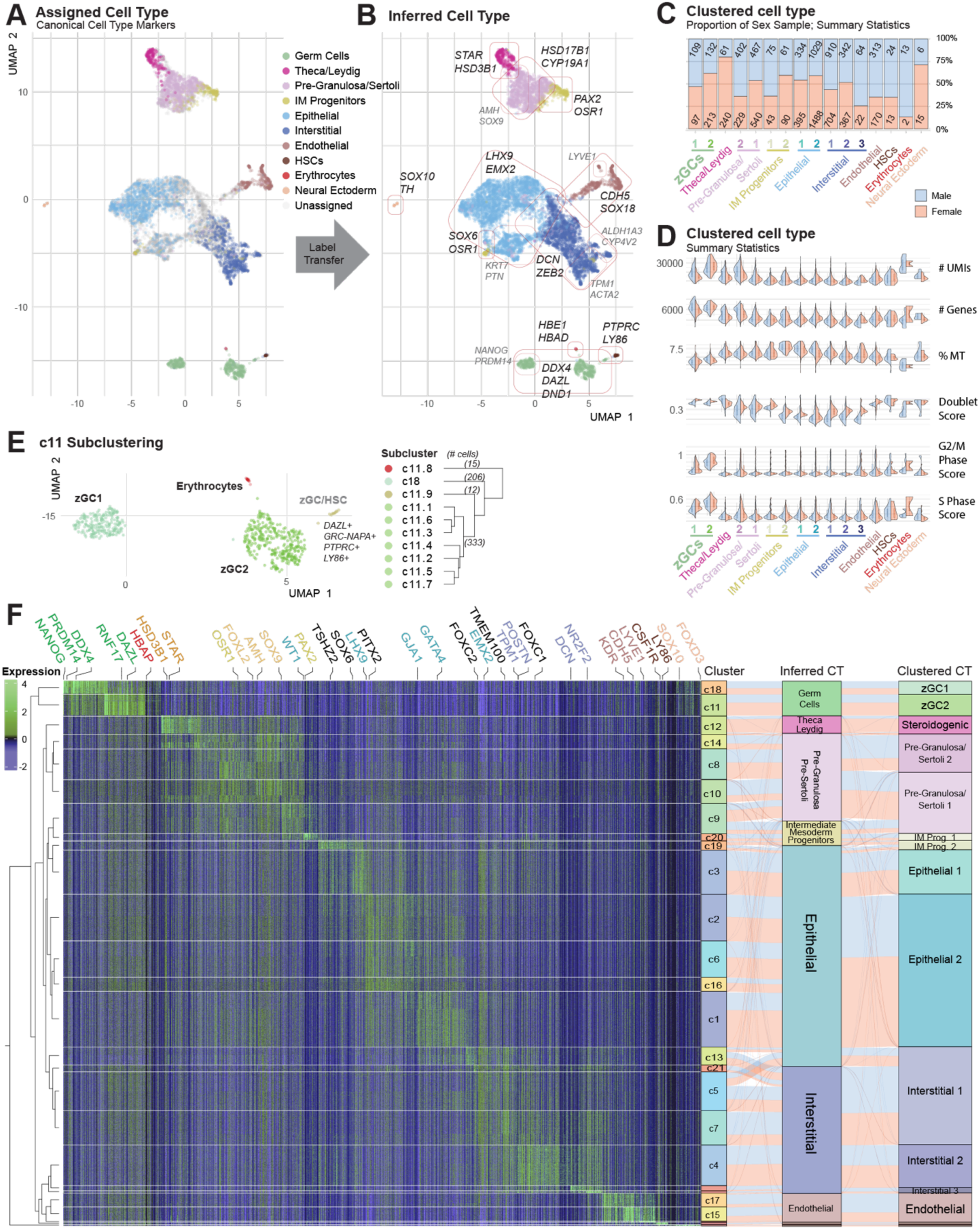
Cell type assessment of HH28 zebra finch gonadal datasets. A. UMAP plot of HH28 zebra finch gonadal cells, colored by assigned cell type using canonical markers (Supplemental table 4). B. UMAP plot of cell type inferred by assigned cell label transfer. C. UMAP of c11 subclustering, with cluster hierarchy dendrogram of subcluster relationships shown on the right. Cluster c11.8 barcodes were redesignated as erythrocytes and c11.1- c11-7 were designated as zGC2. The c11.9 subcluster (n=12) was excluded from subsequent analyses as either an exceedingly rare population not identified by *in situ* hybridization co-labeling (*data not shown*) or as doublet artifacts that had evaded trimming. D. Proportional bar chart of clustered cell types by sex. E. Summary statistics of clustered cell types for each dataset. F. Heatmap of log-normalized expression for gene markers of each cluster, with select gene markers annotated. A dendrogram of hierarchical cluster relationships is provided on the left, and an alluvial plot showing cluster, inferred cell type, and clustered cell type relationships is shown on the right.

**Figure supplement 1.3.**
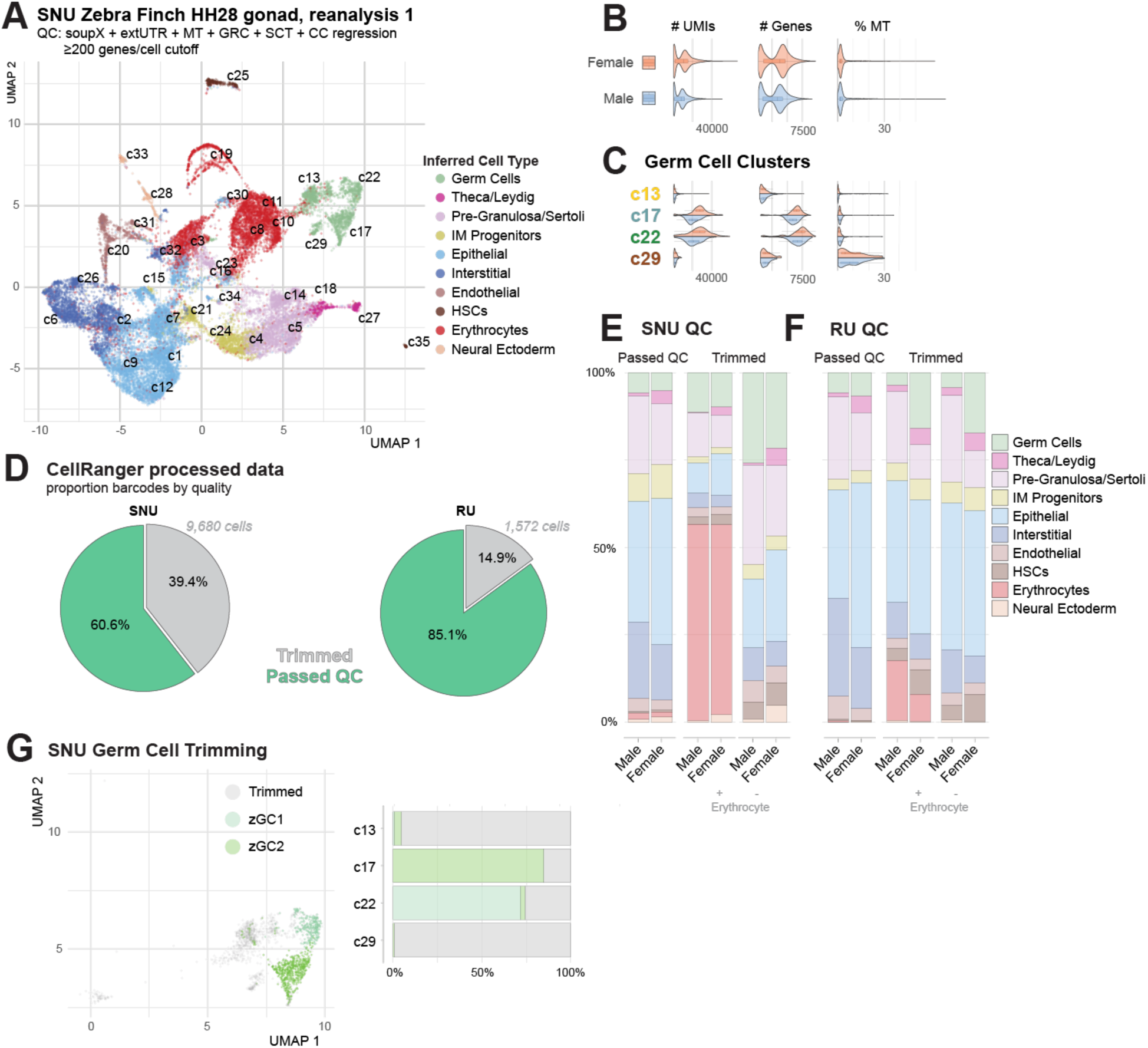
Assessment of SNU zebra finch gonadal datasets. A. UMAP of SNU zebra finch HH28 gonad dataset with analysis methods listed. B. Violin plot of summary statistics for male and female objects. C. Violin plots of summary statistics for each germ cell cluster. D. Pie charts showing proportion of cell barcodes passing additional UMI and Gene count-based quality control measures (Figure 1–figure supplement 1A) in the SNU and RU datasets. E. Inferred cell type demographics of barcodes passing and failing quality control in SNU datasets. F. Inferred cell type demographics of barcodes passing and failing quality control in RU datasets. G. UMAP subset of germ cells, colored by post-quality control identity (left) and corresponding proportional barchart of barcode trimming for each of the initial 4 GC clusters.

**Figure supplement 1.4.**
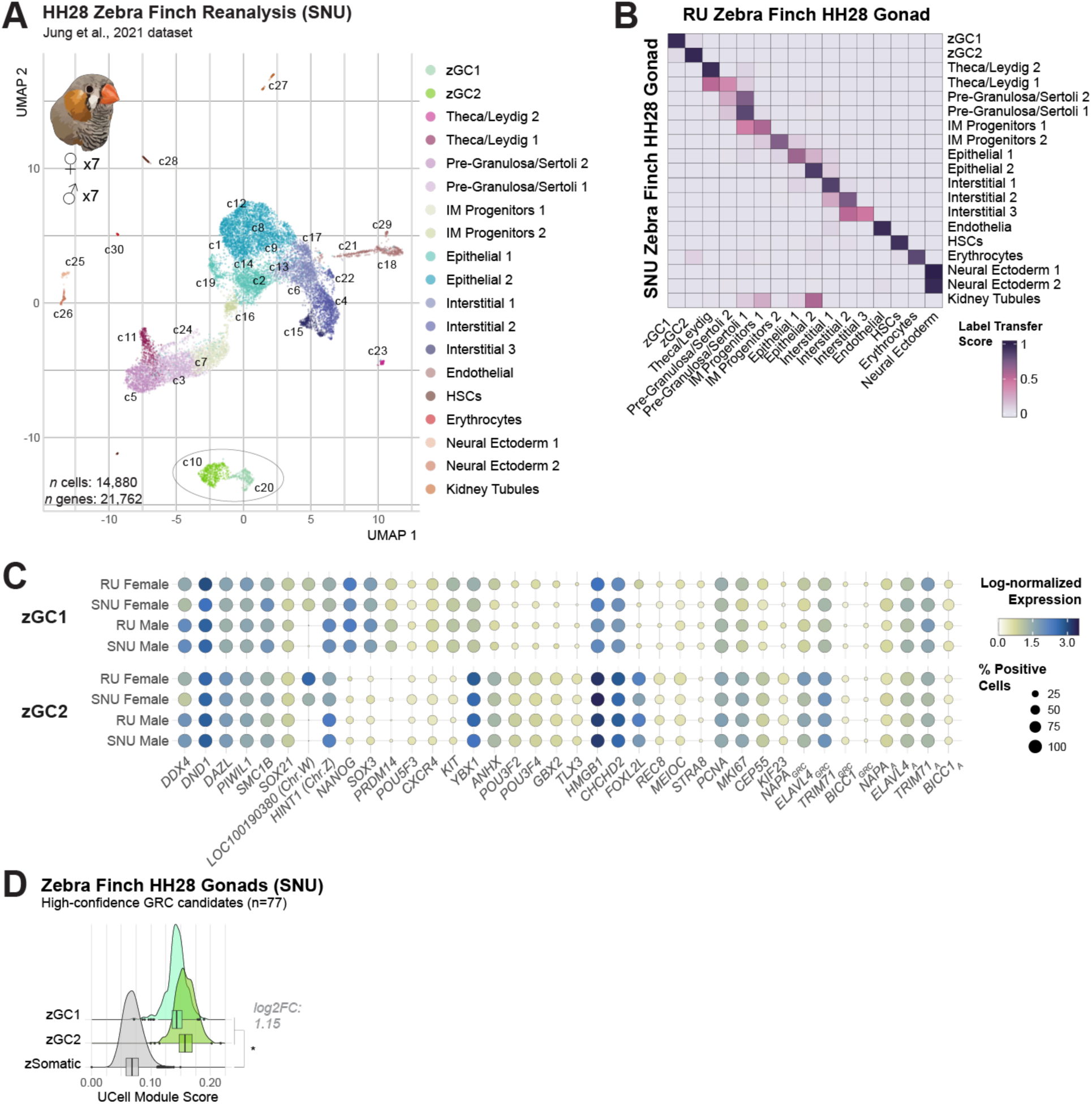
Reanalysis of SNU zebra finch gonadal dataset. A. UMAP plot of reanalyzed single-cell RNAseq datasets from Jung et al., 2021 (SNU), utilizing additional quality control measures. Cells are labeled by independently determined clustered cell types. B. Confusion matrix using Seurat label transfer scores for the SNU dataset and the dataset generated for this study (RU), demonstrating highly concordant gene expression profiles for each cell type in the two datasets. C. Comparative dotplot of log-normalized gene expression between respective zGC1 and zGC2 clusters for each RU and SNU dataset. Note the relative similarity between gene marker expression within clusters. D. Module score comparison between germ cells and somatic cells from the SNU zebra finch datasets. Module is composed of the 77 unmapped, high-confidence zebra finch GRC gene paralog candidates. A log2FC > 0.5 between zGC and zSomatic populations and a p-value ≤ 0.05 by two-sided t-test is denoted by * (Supplemental table 33).

**Figure supplement 2.1.**
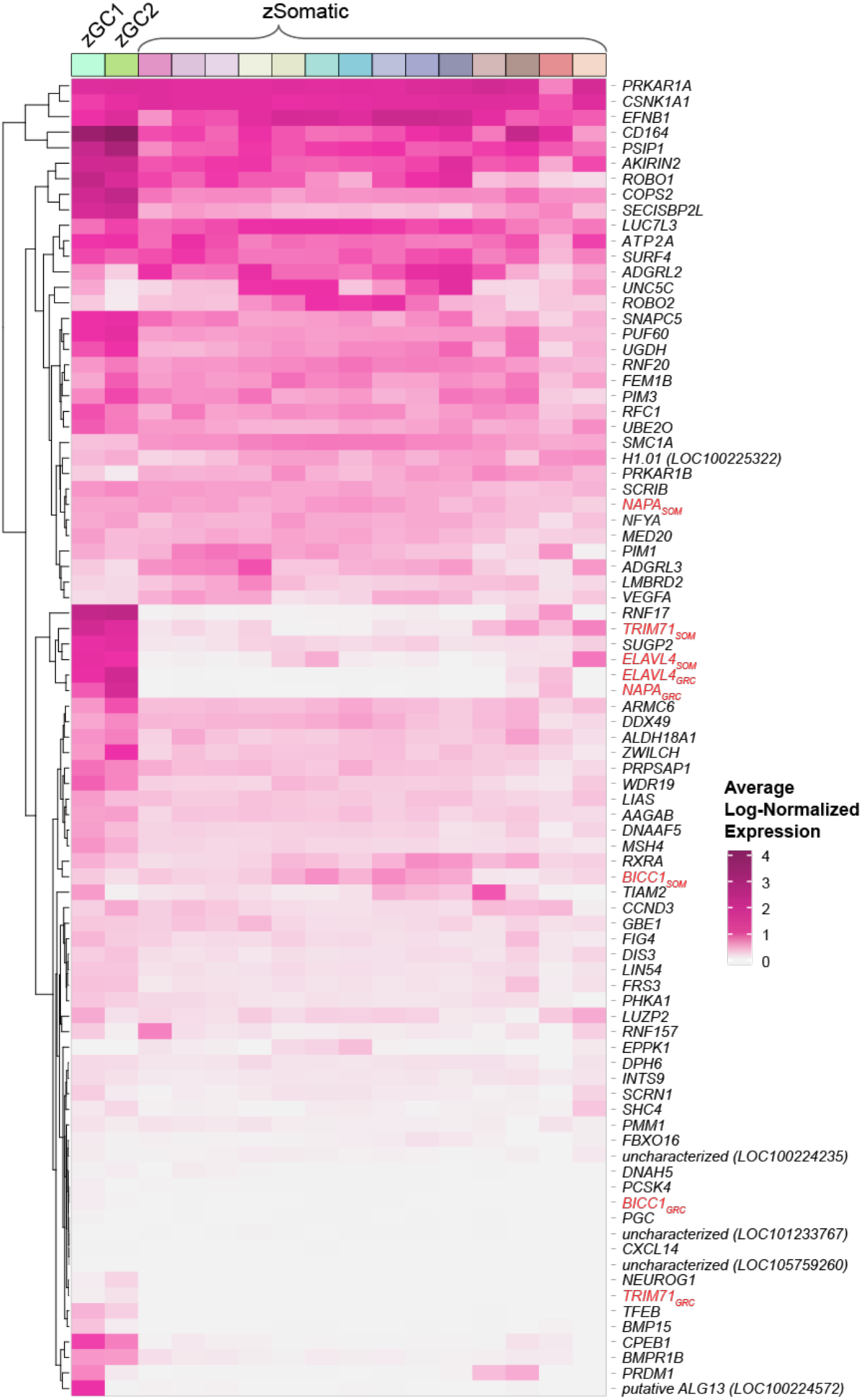
Heatmap of GRC gene candidates. Heatmap showing log-normalized average expression of high confidence GRC paralogs across clustered cell types in zebra finch HH28 data (RU). GRC gene sequences and corresponding somatic chromosome paralogs are highlighted in red.

**Figure supplement 2.2.**
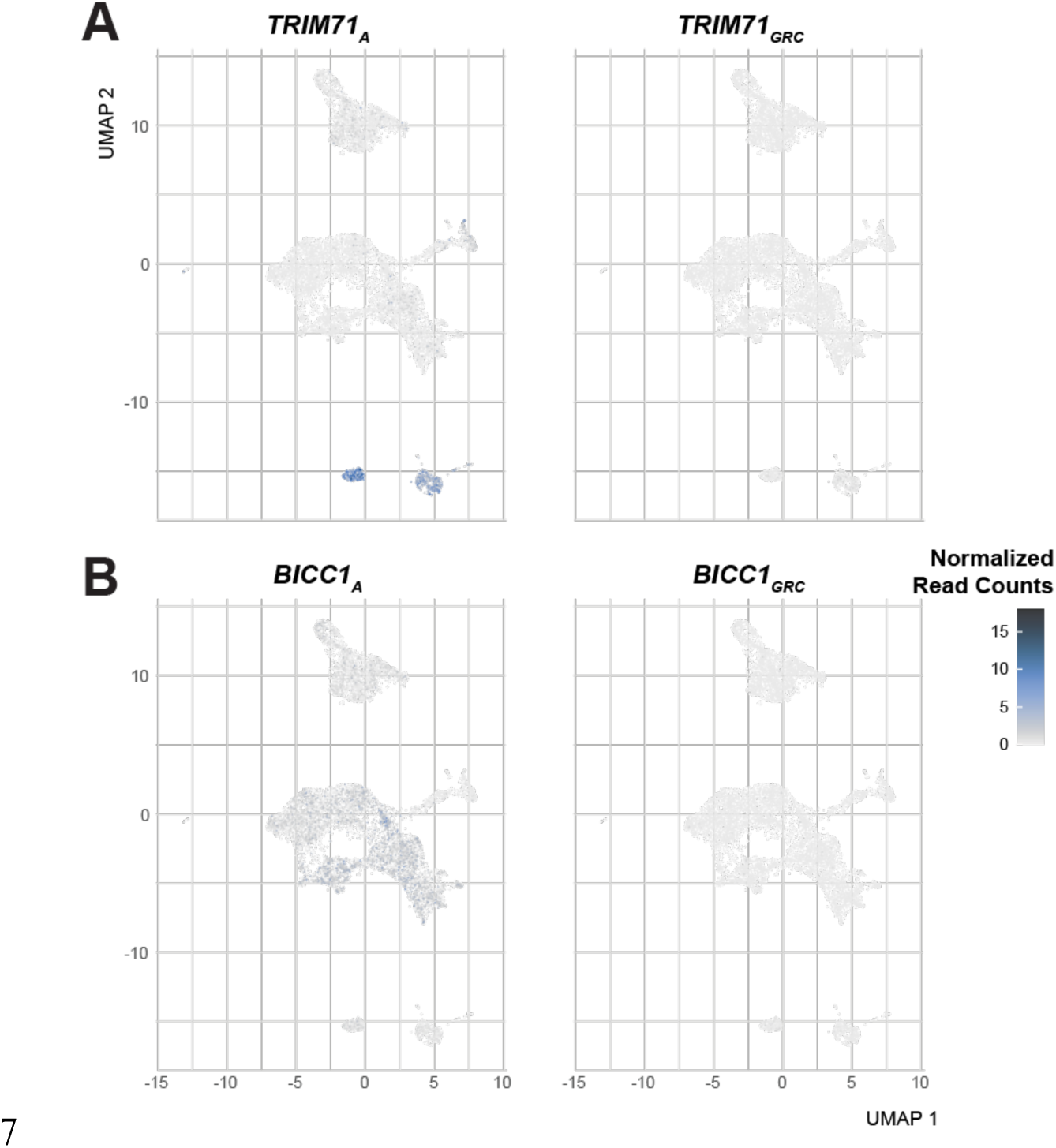
UMAP plots of TRIM71 and BICC1 gene pairs. A. Relative Gene expression counts of TRIM71 gene pairs overlaid on the zebra finch HH28 gonad UMAP plot. B. Relative Gene expression counts of BICC1 gene pairs overlaid on the zebra finch HH28 gonad UMAP plot.

**Figure supplement 2.3.**
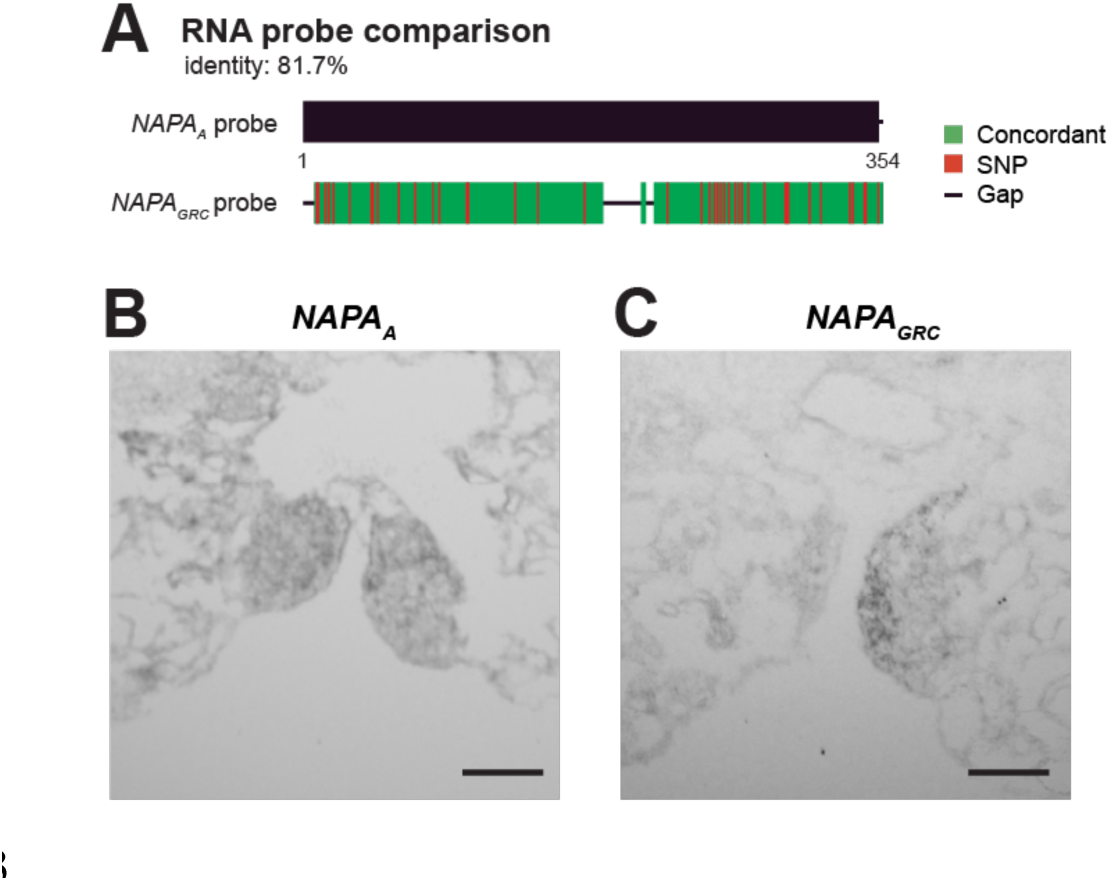
NAPA *in situ* hybridization probe comparisons. A. Illustration of probe sequences, with the *NAPAGRC* probe colored by conservation with the *NAPAA* sequence (basepairs 1-355 of the protein-coding sequence). B. Single-label *in situ* hybridization of embryonic zebra finch gonads with the *NAPAA* probe. C. Single-label *in situ* hybridization of embryonic zebra finch gonads with the *NAPAGRC* probe. Scale bars = 500µm.

**Figure supplement 3.1.**
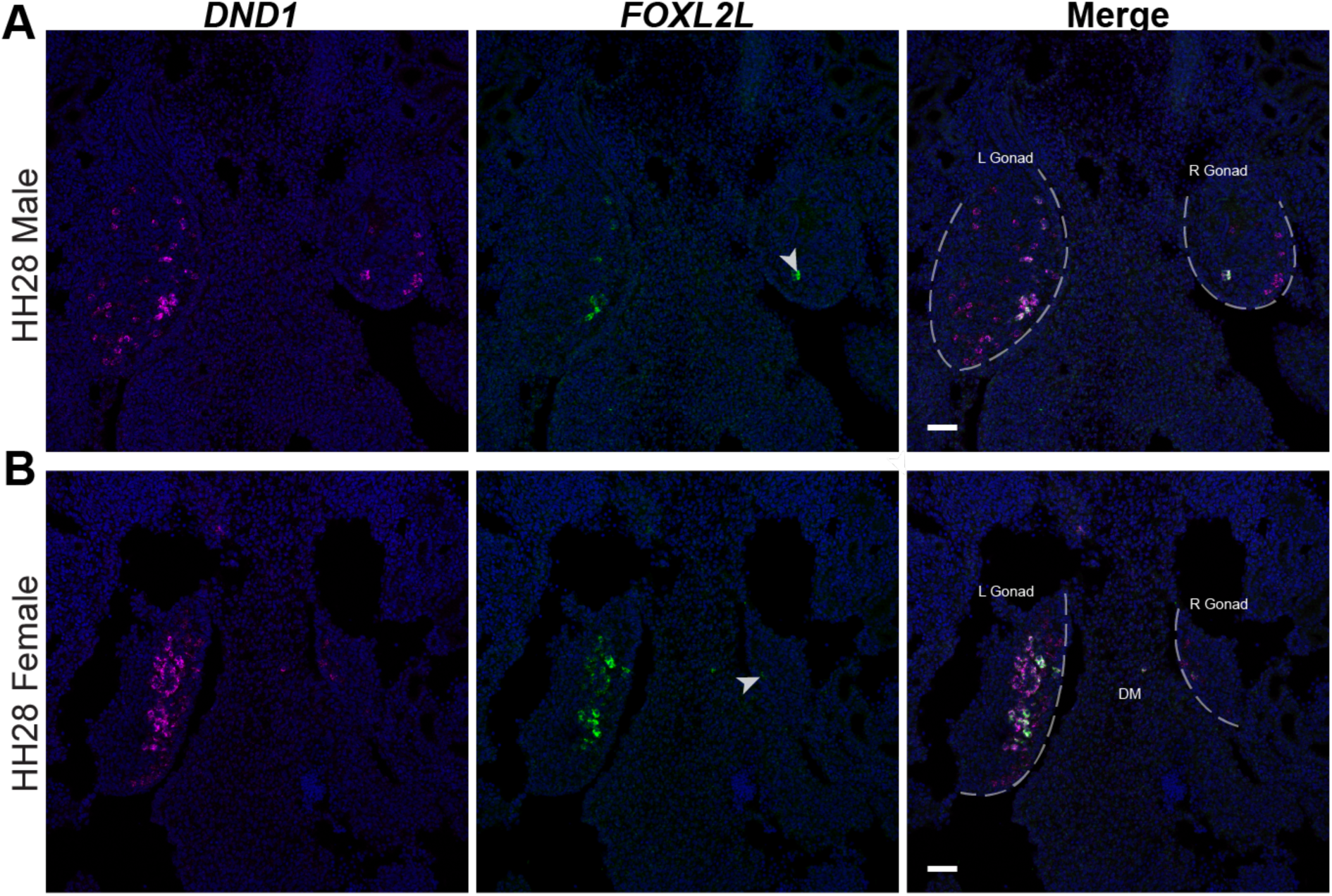
*FOXL2L* expression in the left and right gonad of HH28 zebra finch. A. Dual-label *in situ* hybridization of *DND1* and *FOXL2L* in HH28 male gonads. B. Dual-label *in situ* hybridization of *DND1* and *FOXL2L* in HH28 female gonads. White arrowhead highlights *FOXL2L+* germ cell in the right gonad. Scale bars = 50µm.

**Figure supplement 3.2.**
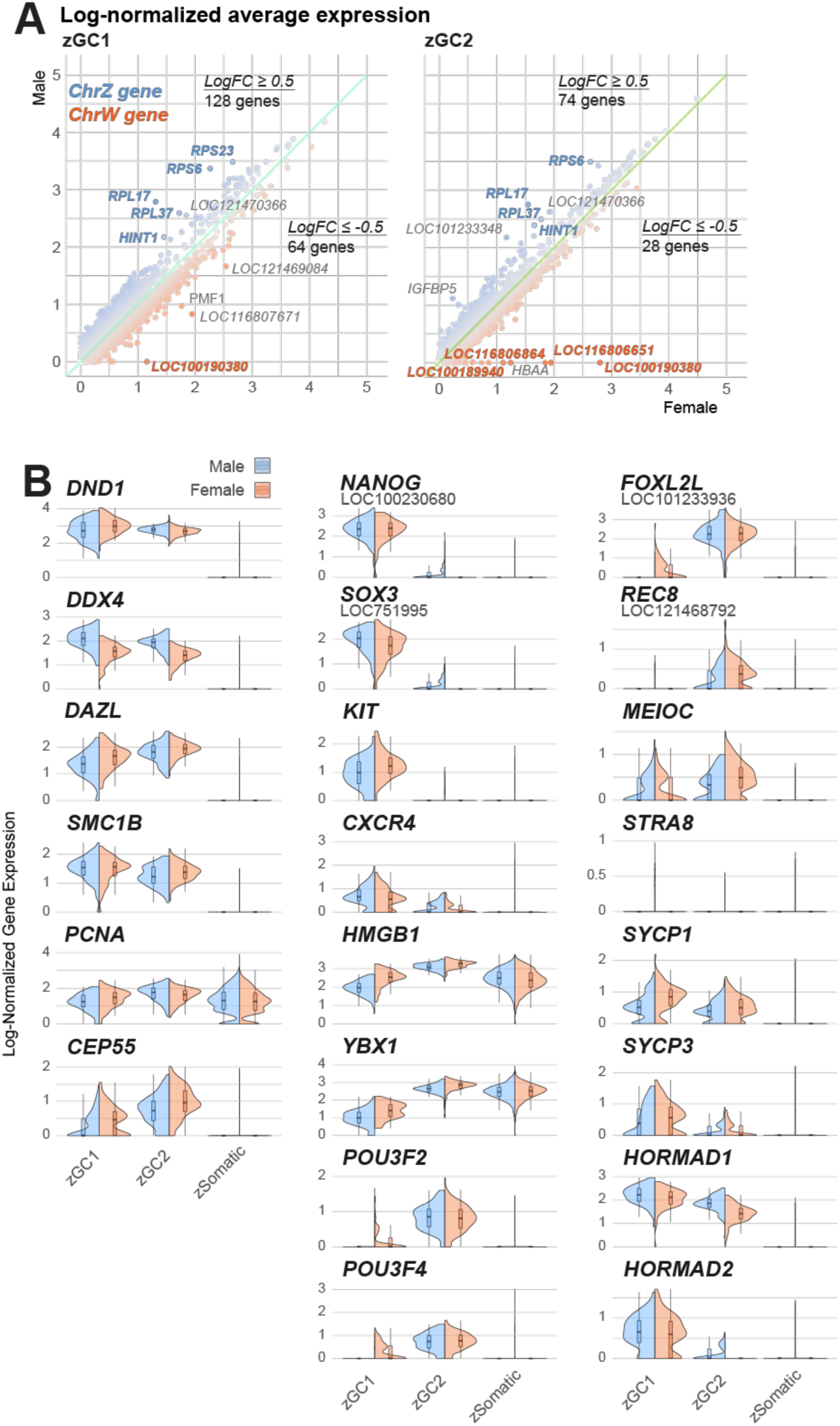
Analysis of zGC sex differences. A. Log-normalized gene expression for zGC1 (left) and zGC2 (right), split by female (x-axis) and male (y-axis) datasets. Points are colored by the relative log-fold change in gene expression between sexes. Highly differential genes located on the sex chromosomes are highlighted in blue (Chr.Z) or red (Chr.W). B. Violin plots of log-normalized gene expression for select genes, split by male (blue) and female (pink) expression.

**Figure supplement 4.1.**
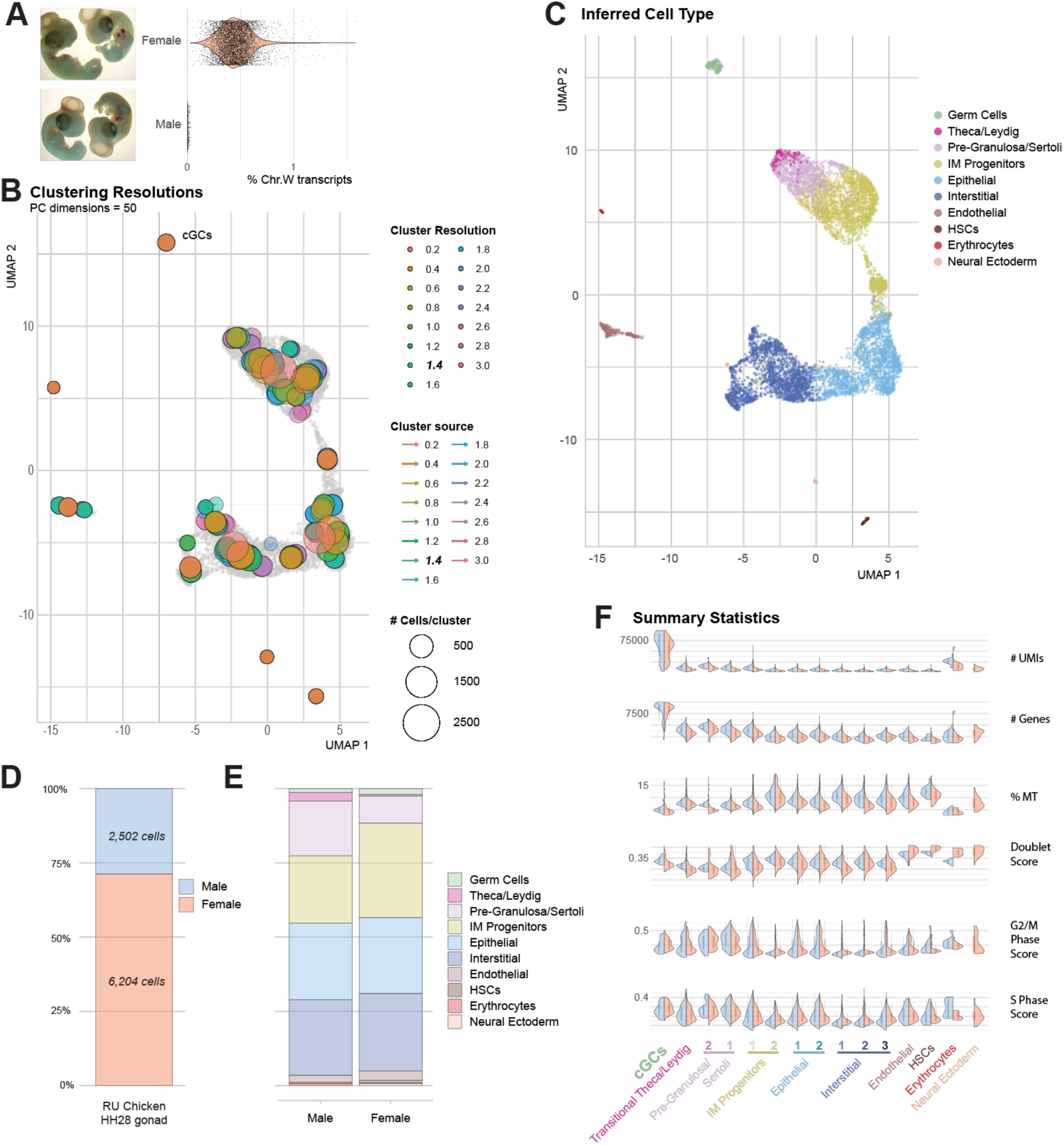
Generation of HH28 chicken gonadal datasets. A. Images (left) of HH28 chicken embryos used to generate female (top) and male (bottom) gonadal datasets. Violin plots (right) validate PCR-based sextyping (not shown) by percentage of W sex chromosome genes expressed. B. UMAP plot overlaid with cluster resolution hierarchy. A clustering resolution of 1.4 was used for subsequent processing. C. UMAP plot showing cell type inferred from cell marker assignments (Supplemental table 3) and subsequent label transfer analysis. D. Proportional bar chart showing difference in total cell number for male and female barcodes. E. Proportional bar chart of inferred cell type for male and female datasets, showing roughly equivalent proportions despite total cell number difference. F. Violin plots of summary statistics, split over clustered cell types.

**Figure supplement 4.2.**
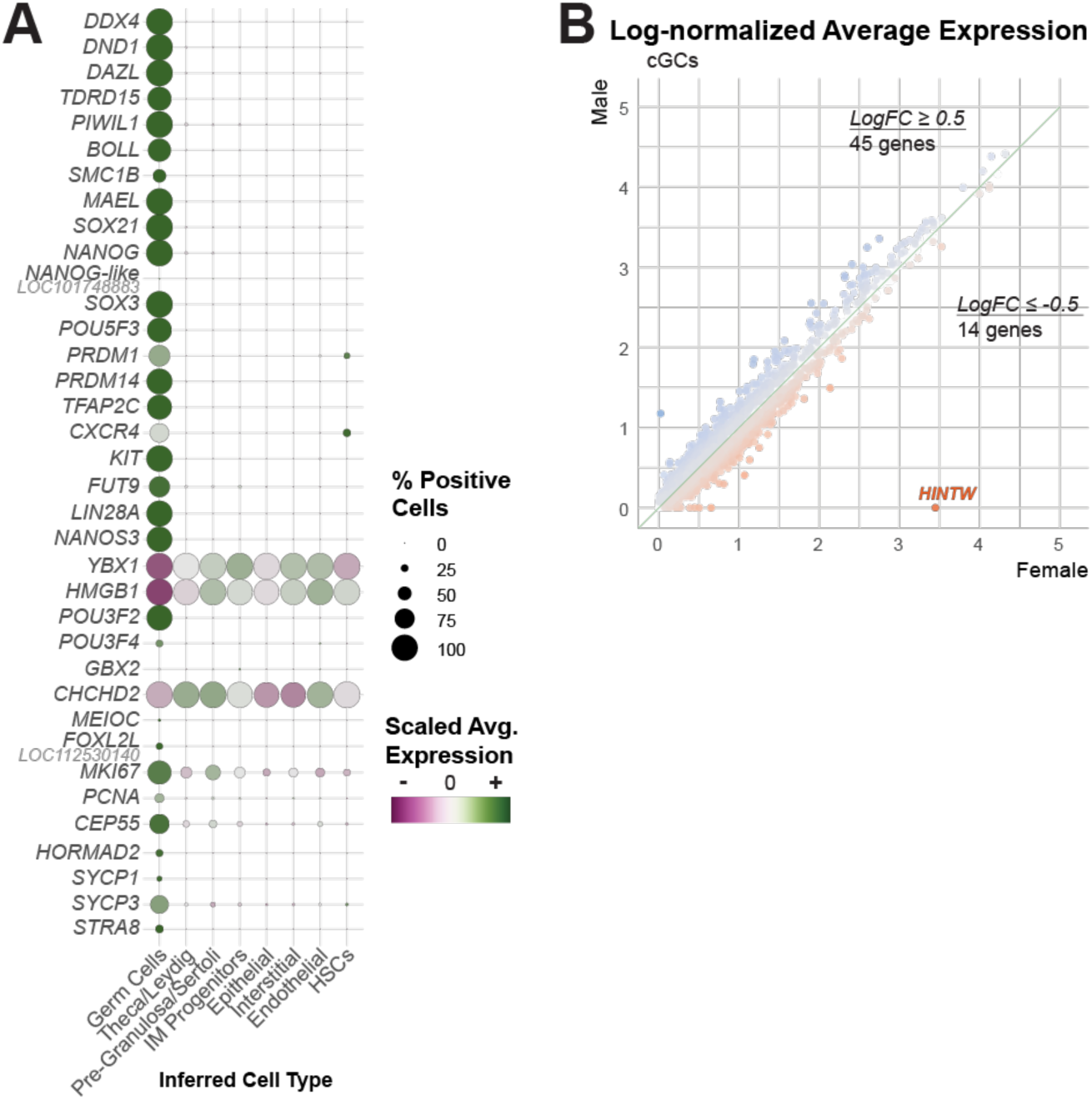
Analysis of cGC gene expression. A. Dot plot of scaled average gene expression of select genes for each inferred cell type in the male and female chicken HH28 gonadal datasets. B. Comparison of average log-normalized gene expression between male and female chicken germ cells (cGCs).

**Figure supplement 4.3.**
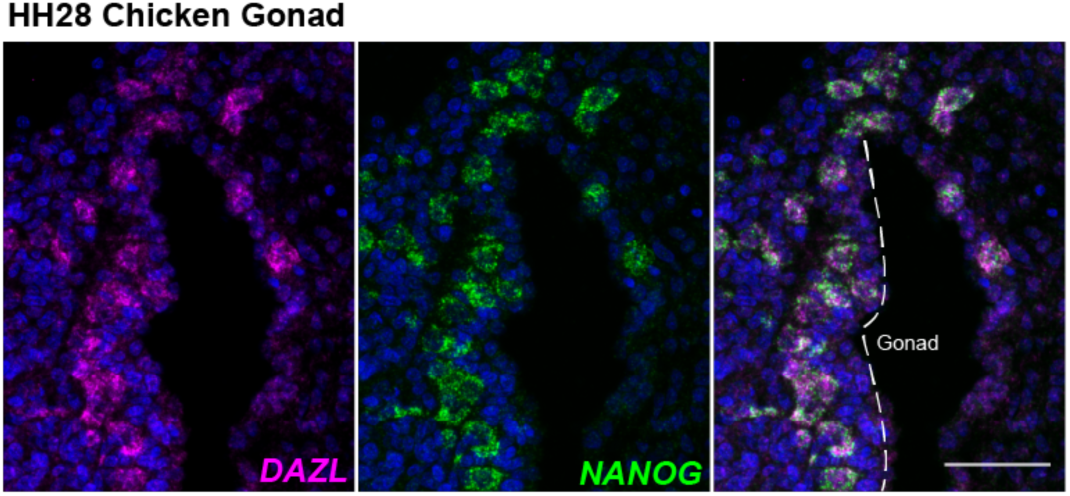
in situ hybridization of *DAZL* and *NANOG* in HH28 chicken gonads. Note the extra-gonadal *DAZL*+/*NANOG*+ cells in the dorsal mesentery, likely migrating into the gonad. Scale bar = 50µm.

**Figure supplement 5.1.**
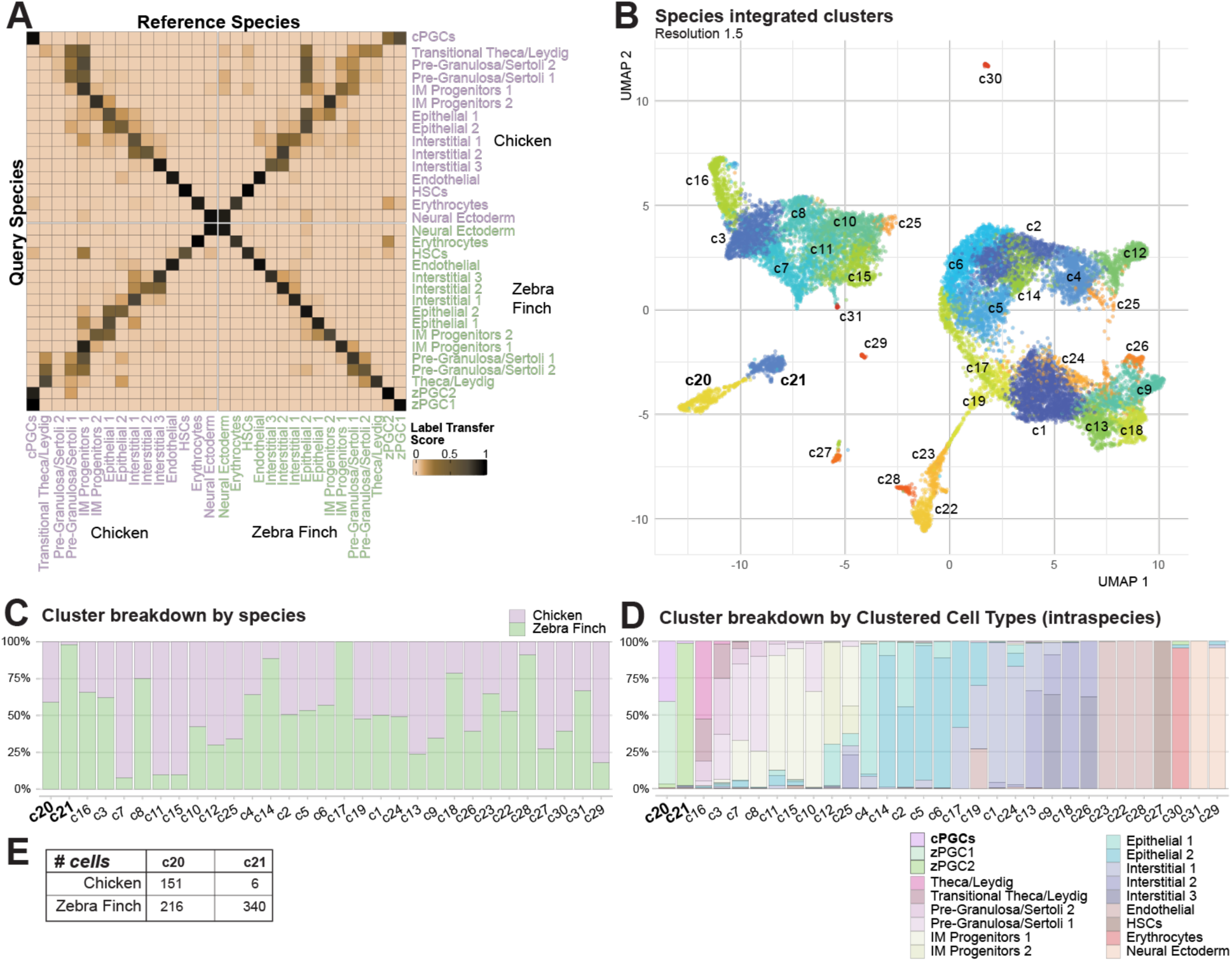
Integrated cross-species dataset analysis. A. Confusion matrix of label transfer similarity scores between clustered cell types of HH28 chicken and zebra finch gonad datasets. B. Integrated UMAP plot of clusters resolved (resolution value = 1.5) for the combined species datasets. C. Proportional bar chart of species contributions for each cluster, with germ cell clusters labeled in bold. Note the high proportion of zebra finch cells in c21. D. Proportional bar chart of inferred cell type contributions for each cluster and specific germ cell clusters highlighted. E. Table demonstrating cell barcode number in each cluster.

**Figure supplement 5.2.**
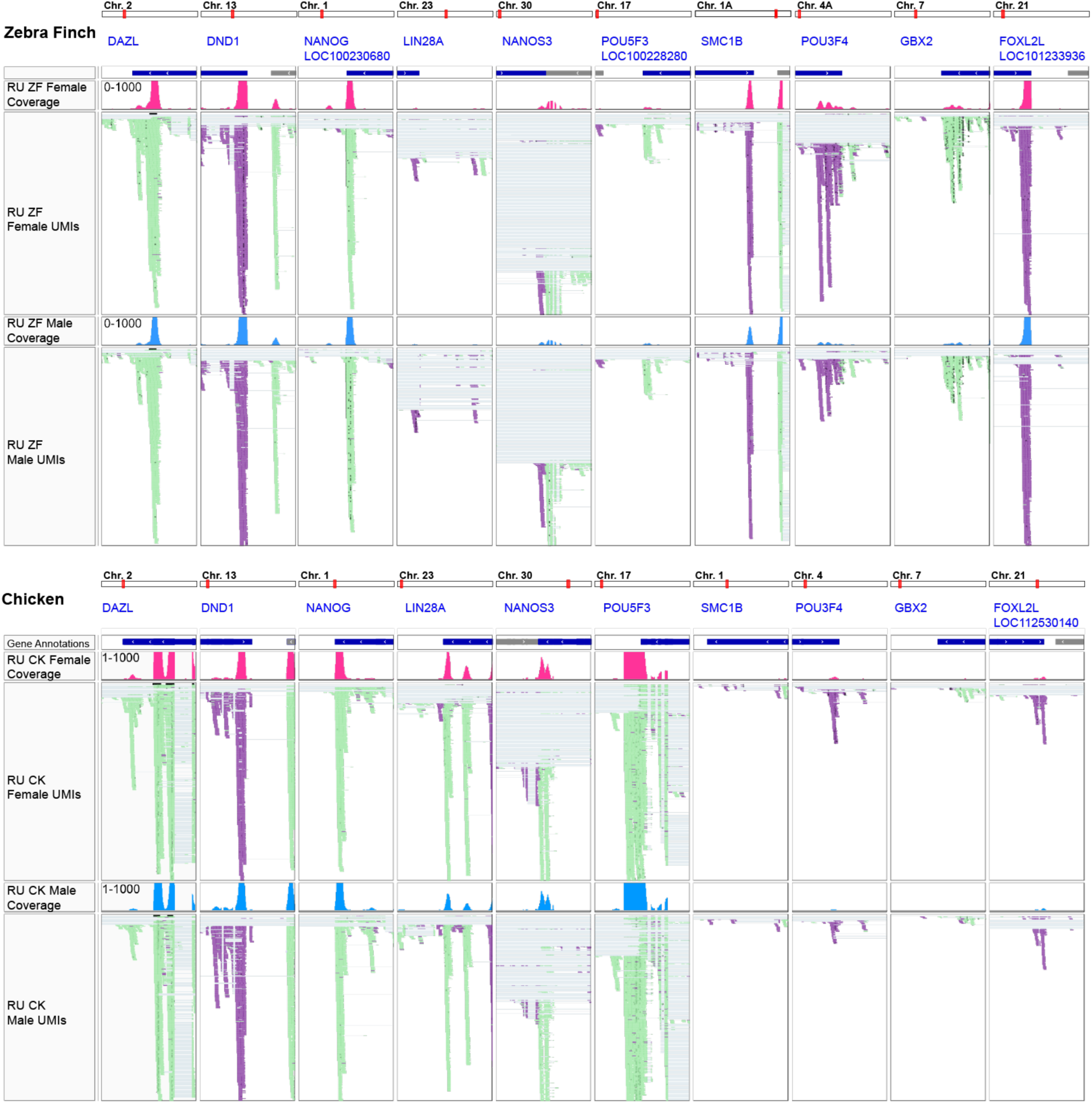
Validation of 3’ gene annotations for differentially expressed orthologs. IGV screenshots of raw read pileups for select orthologs, with minimal read pileups identified outside of the annotations.

**Figure supplement 5.3.**
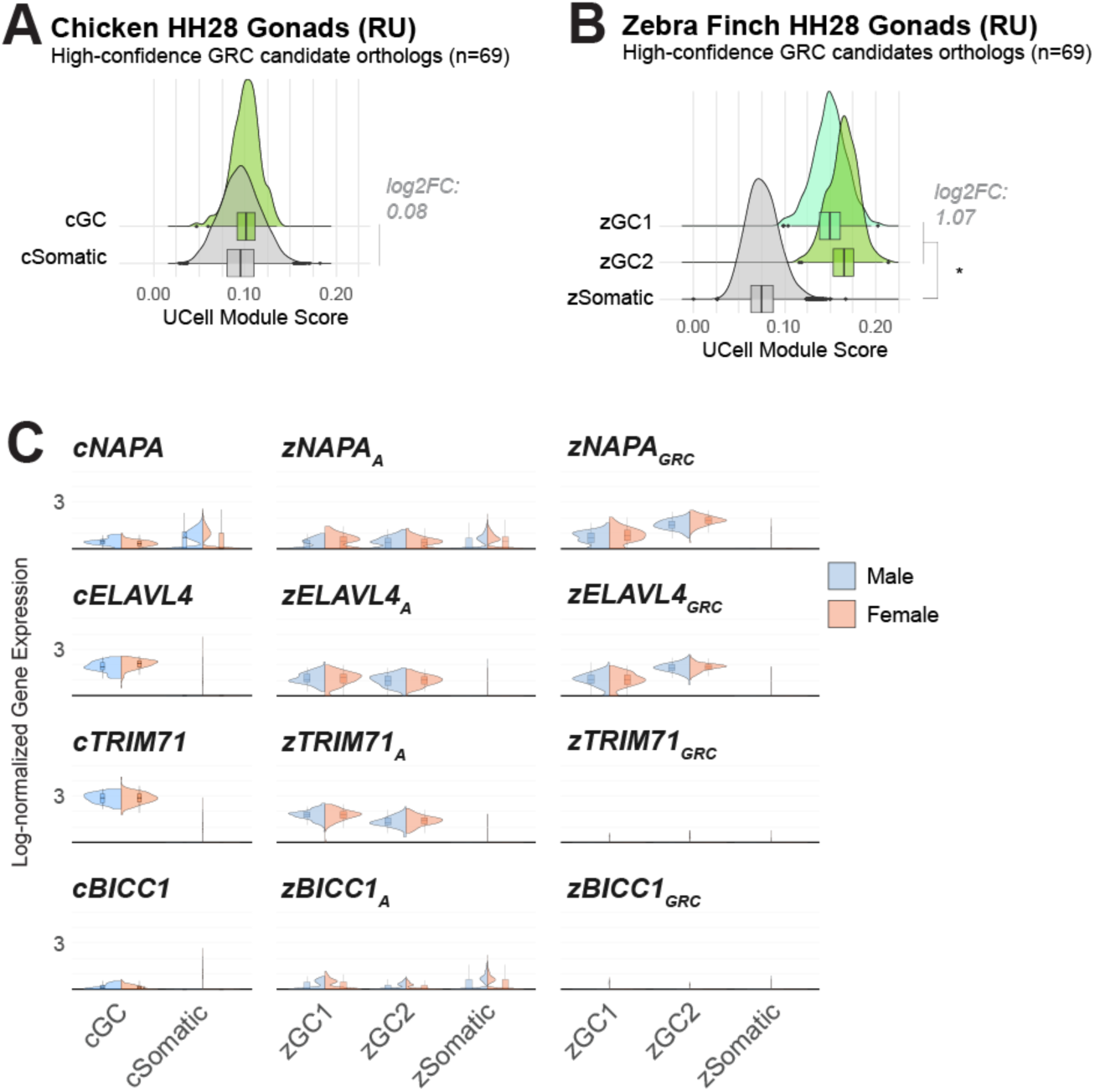
Species comparison of GRC gene candidates. A. Module score comparison between germ cells and somatic cells from the RU chicken datasets. Module is composed of the 69 chicken orthologs of the GRC gene paralog candidates. Log2FC values did not surpass 0.5 between cGC and cSomatic groups. B. Module score comparison between germ cells and somatic cells from the RU zebra finch datasets. Module is composed of the 69 zebra finch GRC gene paralog candidates orthologous to chicken. A log2FC > 0.5 between zGC and zSomatic populations and a p- value ≤ 0.05 by two-sided t-test is denoted by *. C. Violin plots of log-normalized gene expression between male and female cGC and zGC clusters for the each of the four GRC-A gene pairs identified.

**Figure supplement 5.4.**
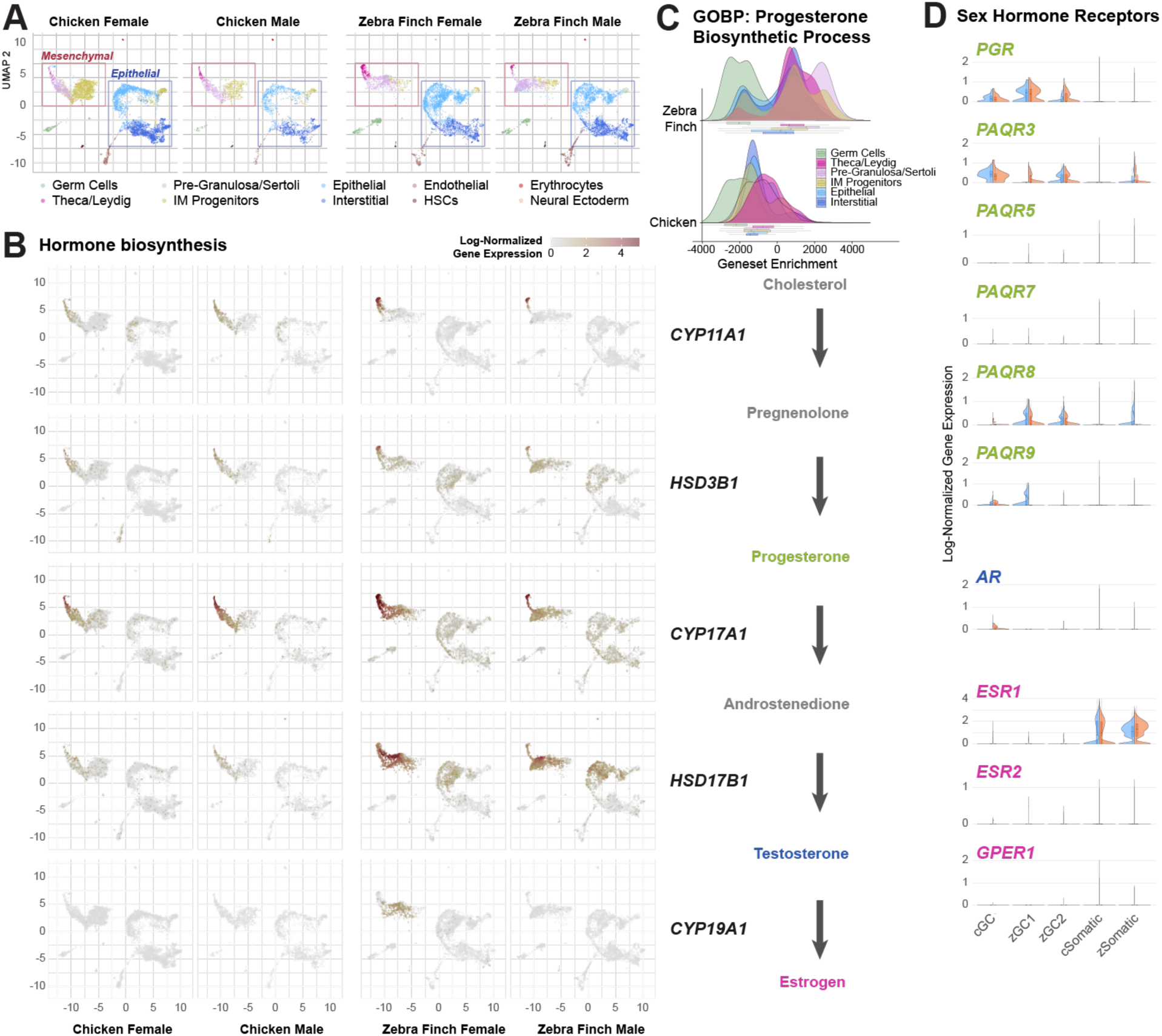
Comparison of sex hormone signaling between chicken and zebra finch datasets. A. Integrated UMAP plots of zebra finch and chicken gonads colored by inferred cell type and separated by dataset. B. Integrated UMAP plots of zebra finch and chicken gonads overlaid with gene markers of retinoid synthesis. C. Ridge plots of progesterone biosynthetic process (GO: 0006701) ssGSEA scores for zebra finch and chicken inferred cell types. D. Violin plots of select sex hormone receptor genes in GC and somatic clusters.

**Figure supplement 5.5.**
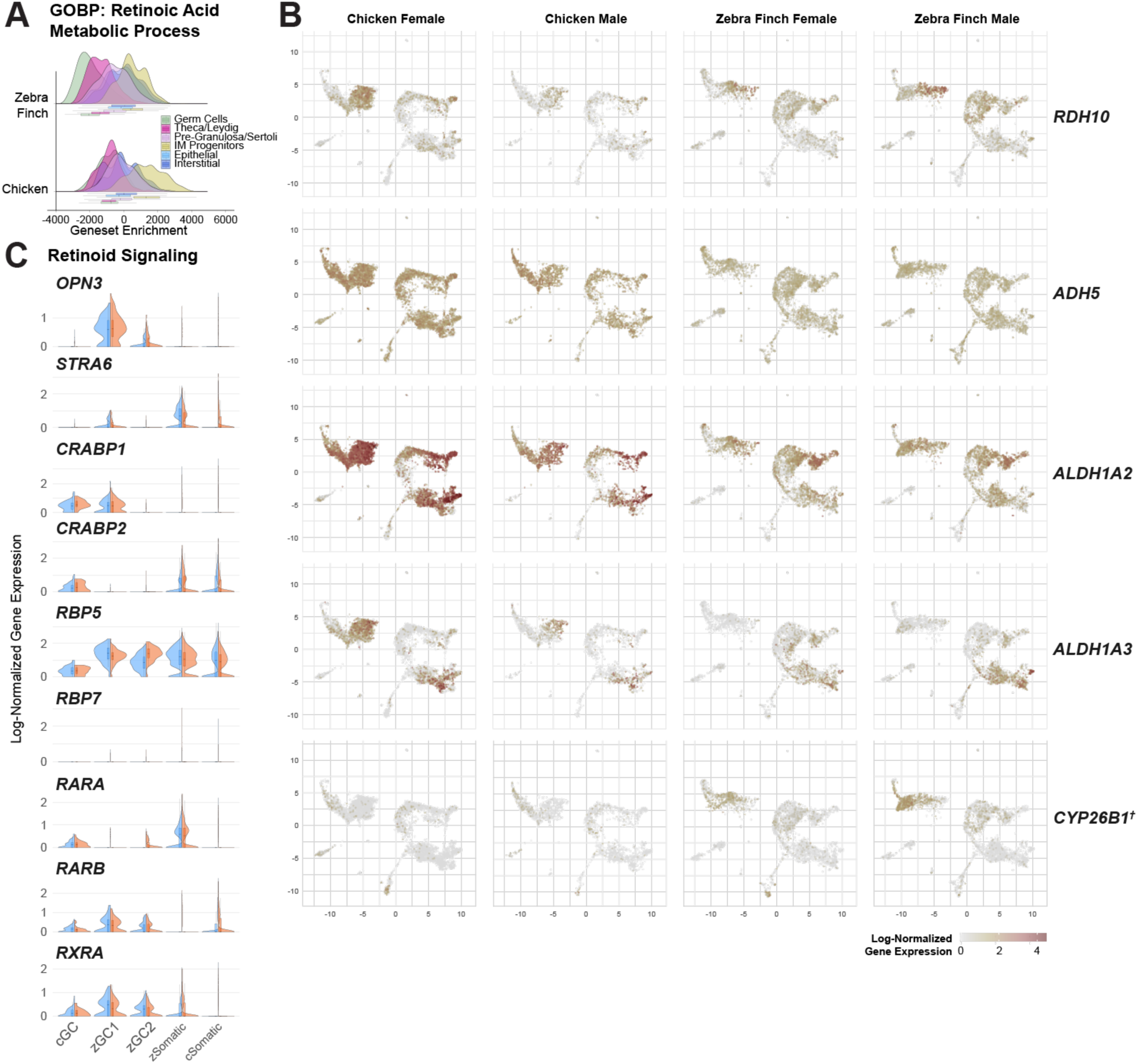
Comparison of retinoic acid signaling between chicken and zebra finch datasets. A. Ridge plots of retinoic acid metabolic process (GO: 0042573) ssGSEA scores for zebra finch and chicken inferred cell types. B. Integrated UMAP plots of zebra finch and chicken gonads overlaid with log-normalized expression for retinoid synthesis and metabolism genes. † denotes chicken and zebra finch *CYP26B1* (LOC100221095) annotations as not formally assigned as ortholog pairs. C. Violin plots of select genes related to retinoid signaling in GC and somatic clusters.

**Figure supplement 6.1.**
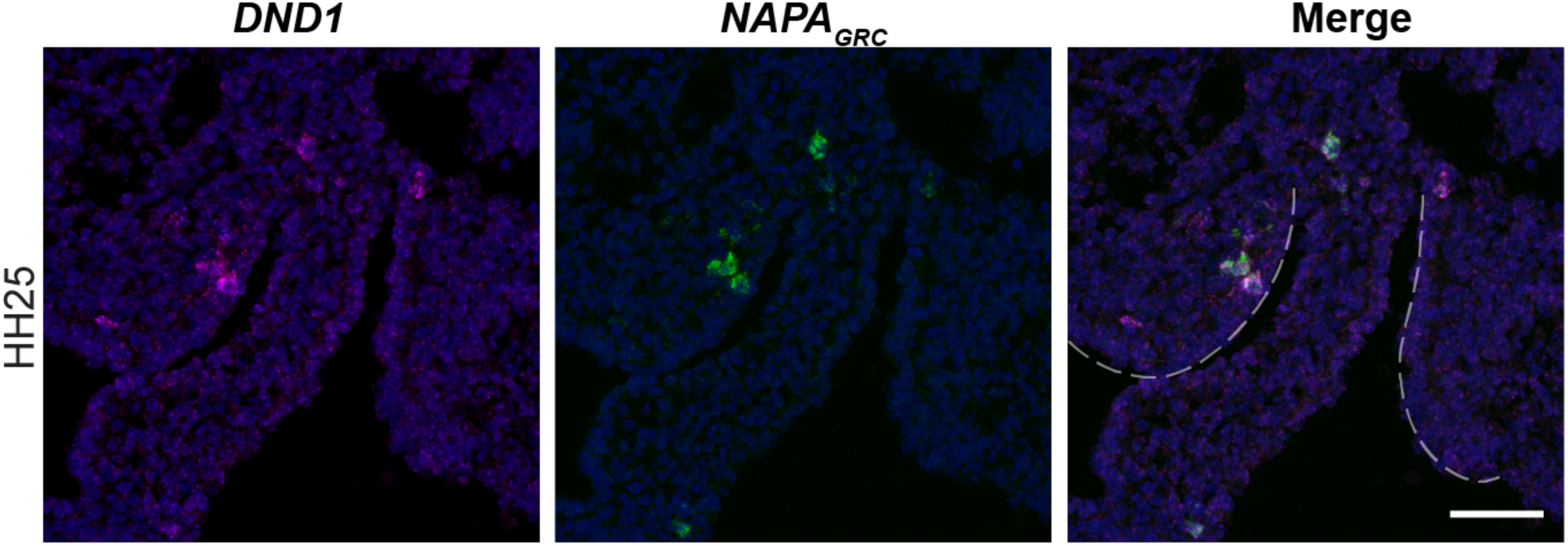
*NANOG* expression in the left and right gonad of HH25 zebra finch. Dual-label *in situ* hybridization of *DND1* and *NANOG* in HH25 male gonads. White dotted lines denote medial gonadal boundaries. Scale bar = 50µm.

**Figure supplement 6.2.**
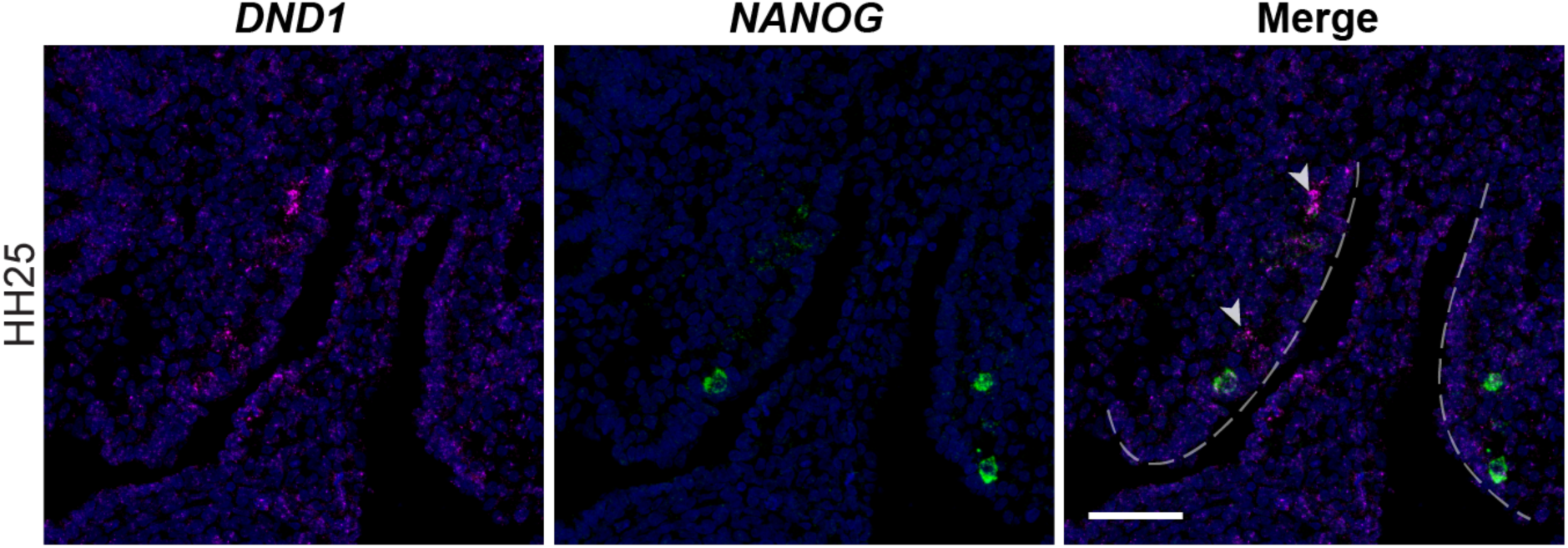
*NAPAGRC* expression in the left and right gonad of HH25 zebra finch. Dual-label *in situ* hybridization of *DND1* and *NAPAGRC* in HH25 male gonads. White dotted lines denote medial gonadal boundaries. Scale bar = 50µm.

**Figure supplement 7.1.**
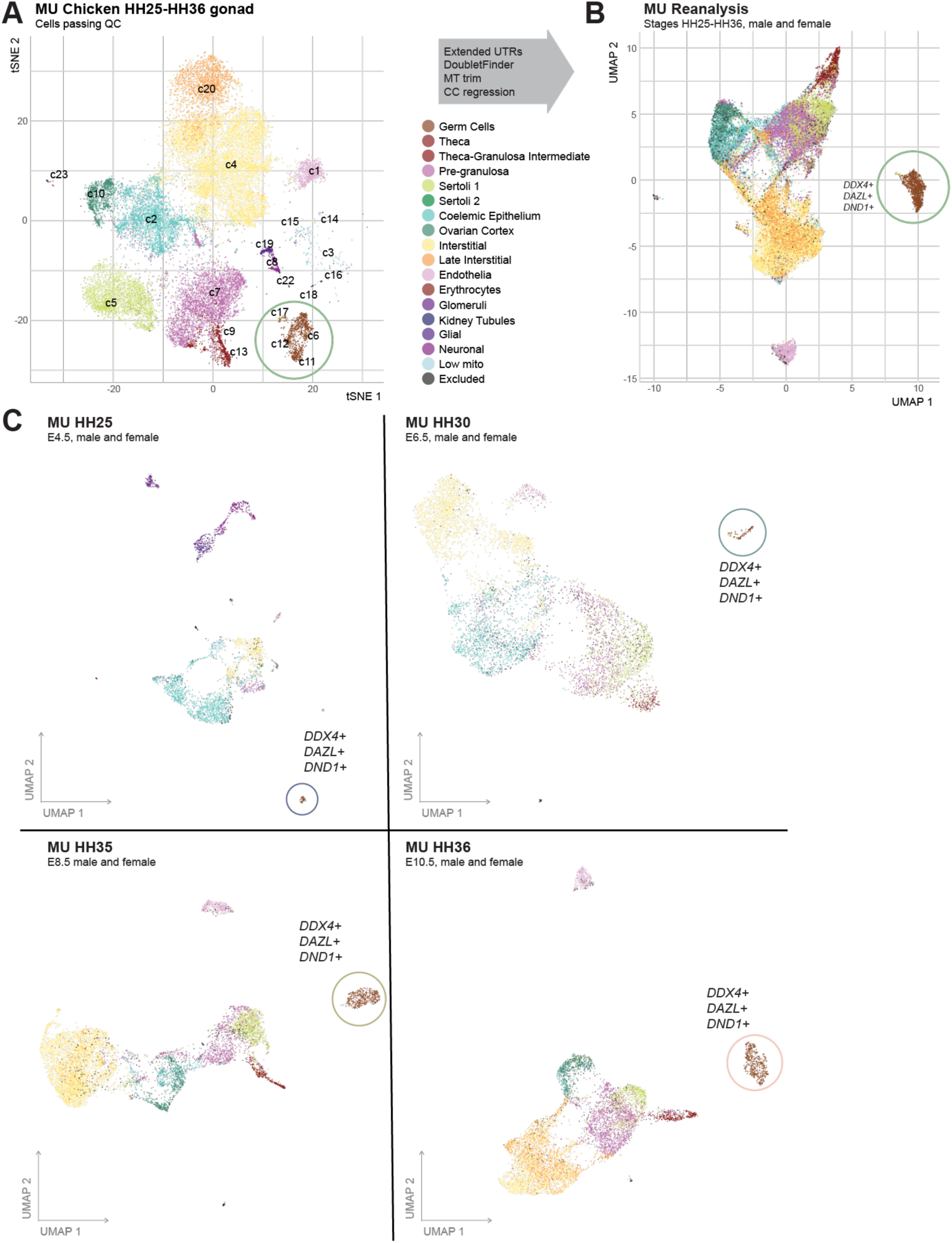
Estermann et al., 2020 dataset (MU) reanalysis. A. Abridged t-SNE projection from Estermann et al., 2020. Barcodes that did not meet quality control measures used for this study were excluded from this plot. Germ cell clusters are circled in green. B. UMAP plot of re-analyzed cell barcodes, including barcodes not assessed in Estermann et al., 2020 that met the quality control measures used in this study (black). Germ cell clusters are circled in green. C. UMAP plots of datasets by embryonic stage. Germ cell clusters are circled for each stage.

**Figure supplement 7.2.**
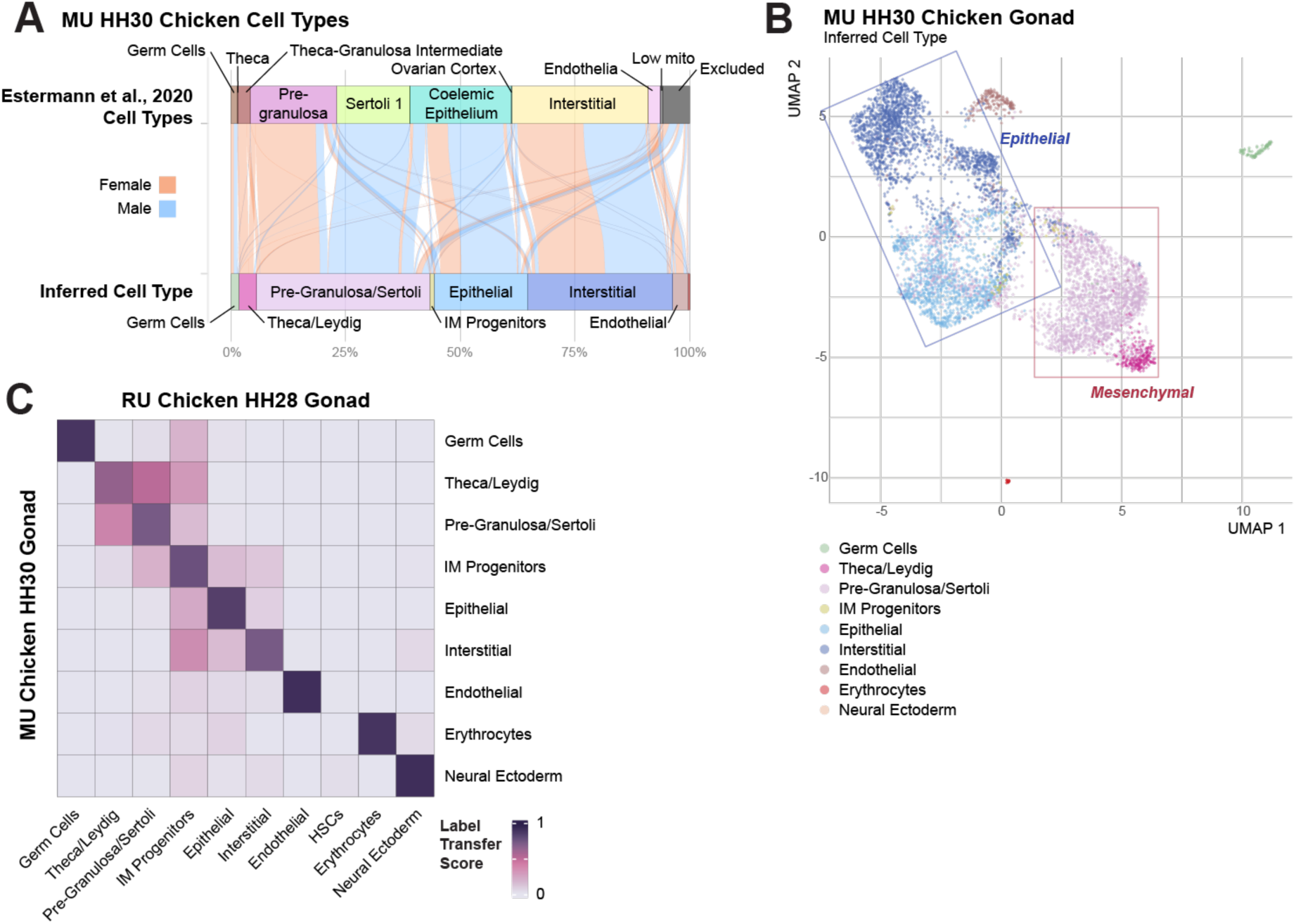
Comparison of Chicken HH28 (RU) and HH30 (MU) gonadal samples. A. Alluvial plot showing relationship between Estermann et al., 2020 HH30 chicken gonad cell types and cell types inferred by label transfer analysis by cell marker assignment. B. UMAP plot showing inferred cell types for MU HH30 datasets. C. Confusion matrix of label transfer similarity scores between inferred cell types of the chicken HH28 (RU) and HH30 (MU) gonadal datasets.

**Figure supplement 7.3.**
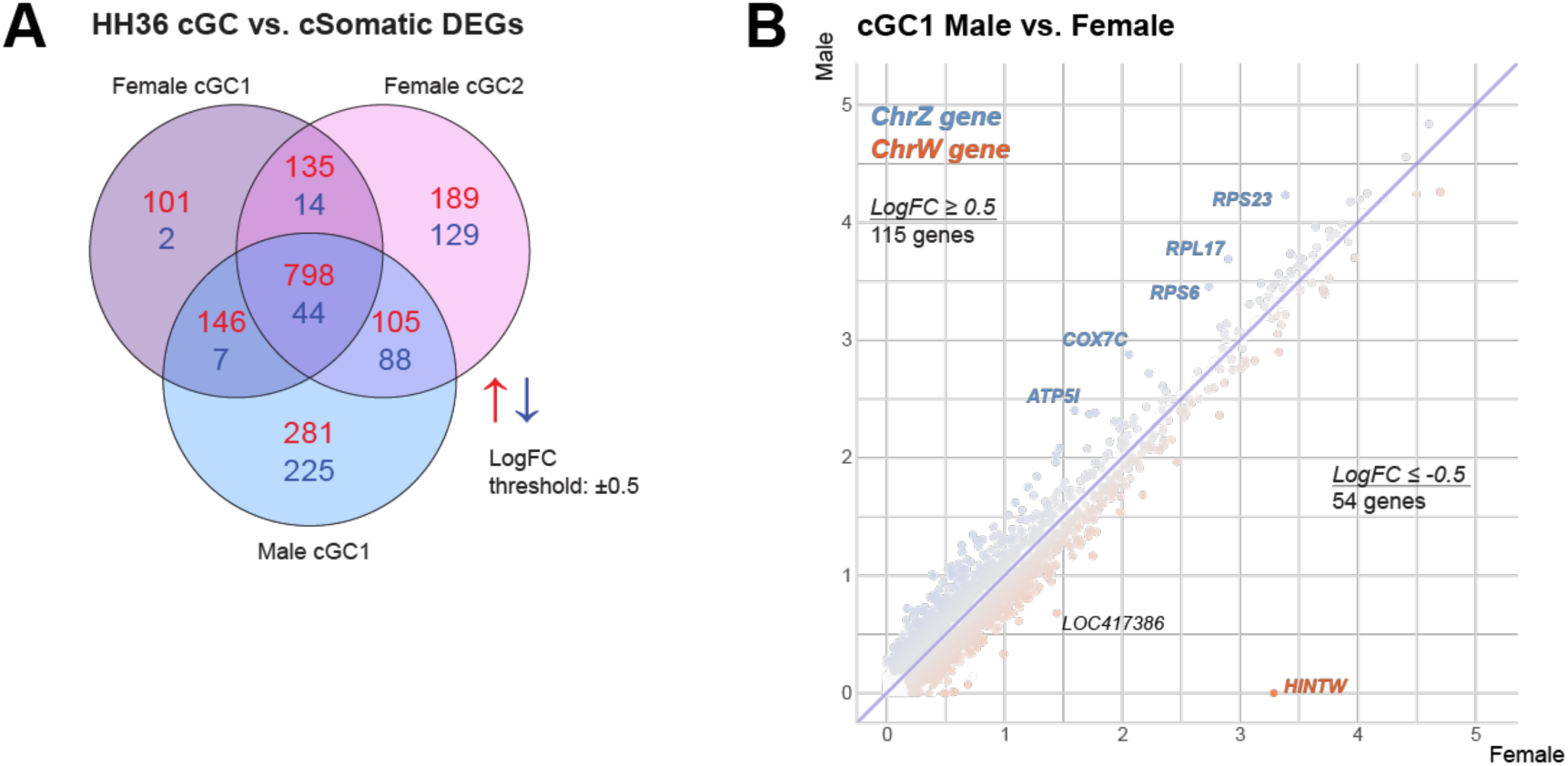
Extended analysis of MU HH36 cGC clusters. A. Venn diagram of upregulated (red) and downregulated (blue) gene expression between male and female cGC clusters and their sex-respective somatic cell types. A differential expression threshold is defined at a log-fold change of ±0.5. B. Log-normalized gene expression of Male (y-axis) and Female (x-axis) cGC1 clusters for each gene. Points are colored by the relative log-fold change in gene expression between clusters, with gene symbol labels for the most differential genes. Label colors denote Chr.W (red) or Chr.Z (blue) gene location.

**Figure supplement 7.4.**
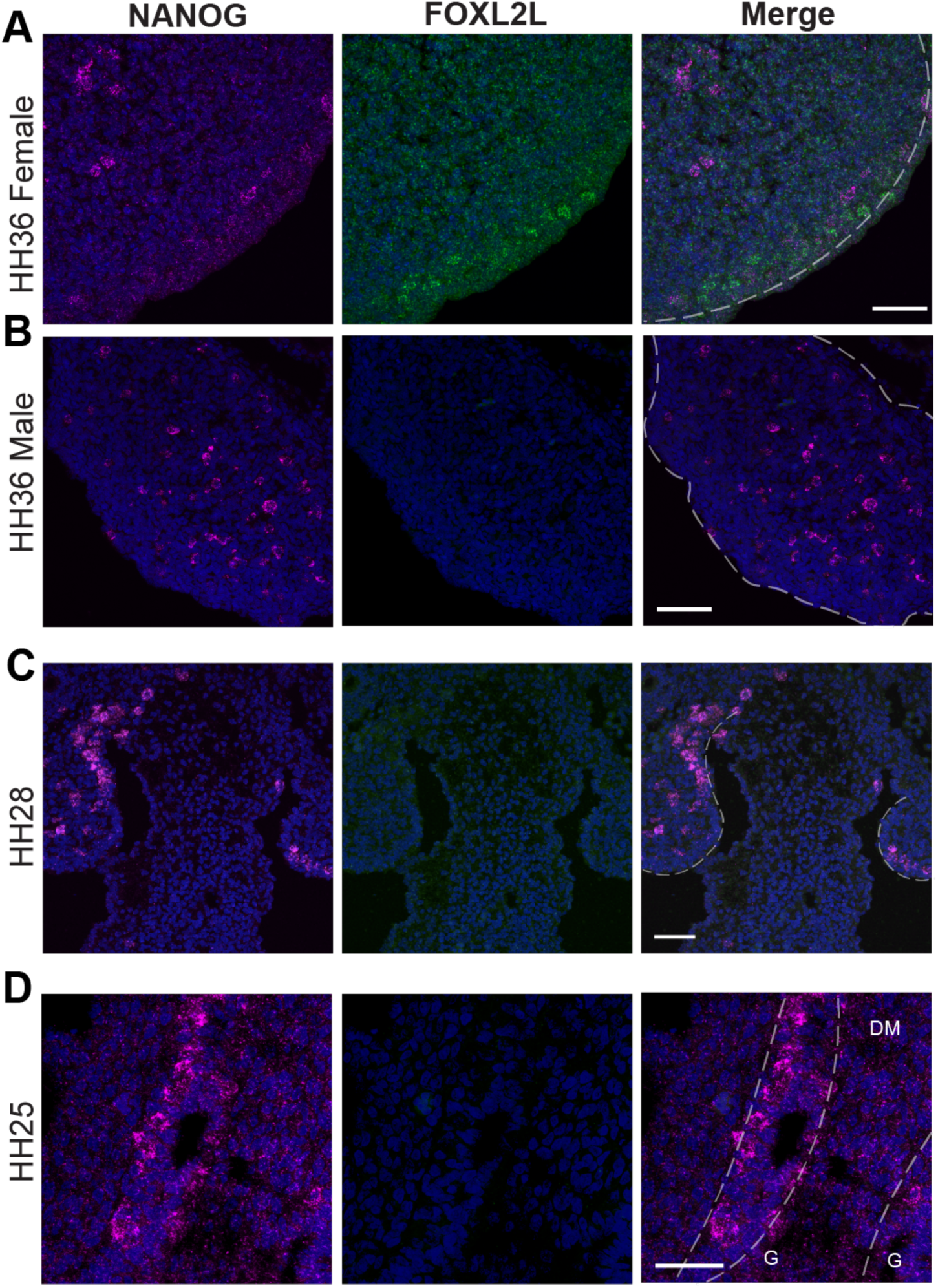
in situ hybridization of *NANOG* and *FOXL2L* across chicken gonadal development. A. Dual-label *in situ* hybridization of *NANOG* and *FOXL2L* in HH36 female chicken gonads. B. Dual-label *in situ* hybridization of *NANOG* and *FOXL2L* in HH36 male chicken gonads. C. Dual-label *in situ* hybridization of *NANOG* and *FOXL2L* in HH28 bipotential chicken gonads. D. Dual-label *in situ* hybridization of *NANOG* and *FOXL2L* in HH25 bipotential chicken gonads. Scale bars = 50µm. G = gonads; DM = dorsal mesentery.

**Figure supplement 7.5.**
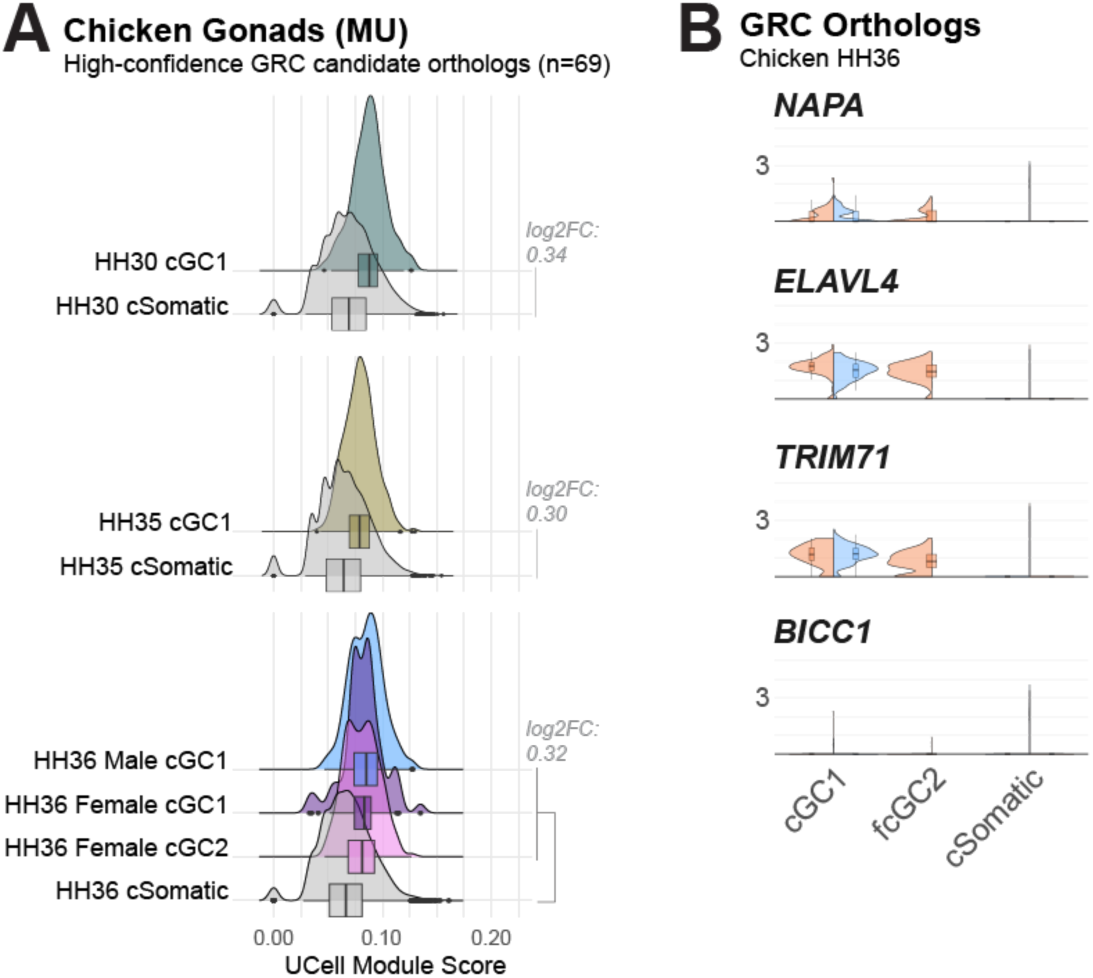
Analysis of GRC ortholog expression in cGC clusters. A. Module score comparison between germ cells and somatic cells for each stage of the MU chicken datasets. Module is composed of the 69 chicken orthologs of the GRC gene paralog candidates. Log2FC values did not surpass 0.5 between cGC and cSomatic groups. B. Violin plots of log-normalized gene expression between male and female HH36 cGC clusters for the each of the four GRC-A gene pairs identified.

**Figure supplement 7.6.**
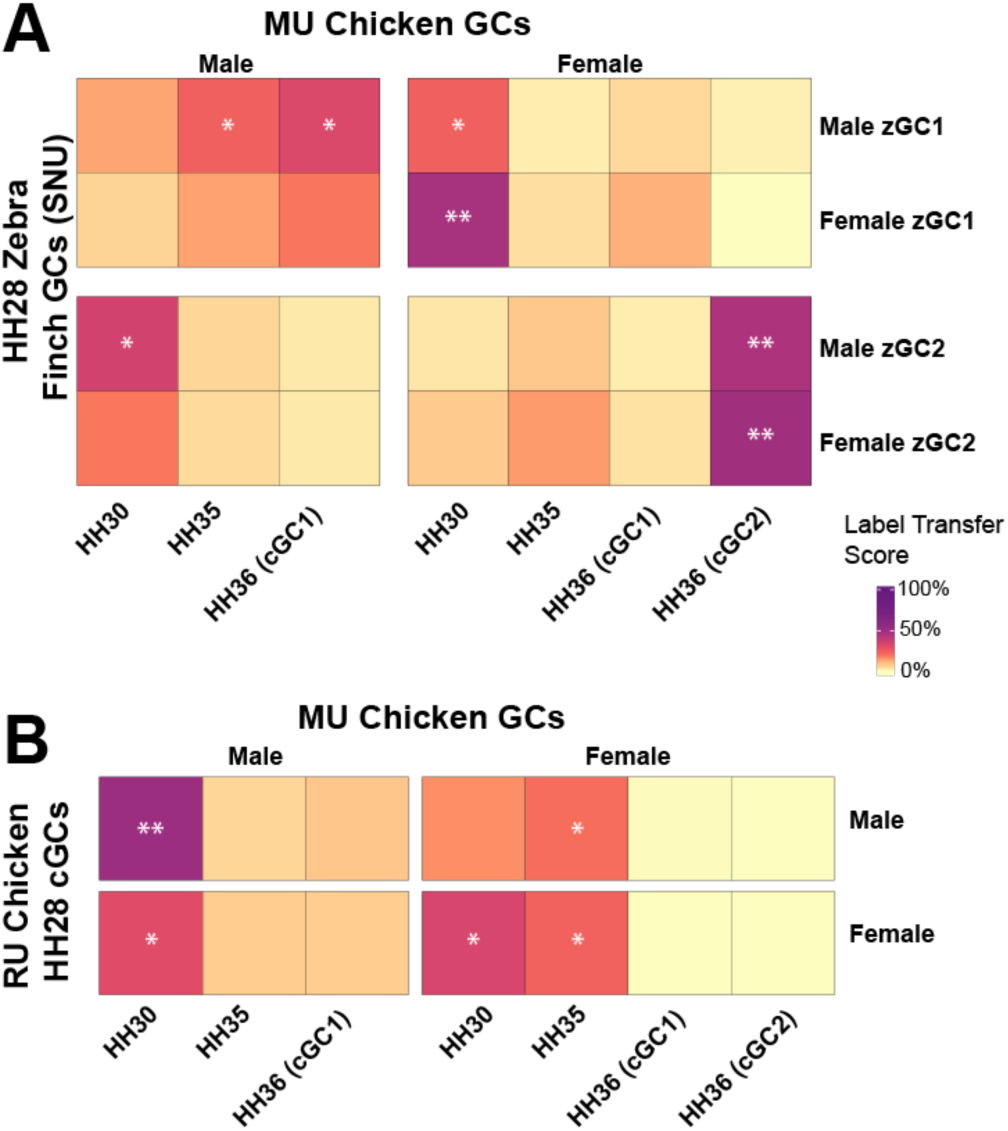
Mapping of additional datasets onto MU cGC expression profiles. A. Confusion matrix of label transfer similarity scores for male and female zebra finch zGC clusters (SNU) against chicken (MU) germ cells at HH30, HH35, and HH36. A log2FC>0.50 against aggregate of scores and a p-value≤0.05 by one-sided t-test is denoted by *. A log2FC>2.0 is denoted by **. B. Confusion matrix of label transfer similarity scores for male and female chicken cGC cluster (RU) against chicken (MU) germ cells at HH30, HH35, and HH36. A log2FC>0.50 against aggregate of scores and a p-value≤0.05 by one-sided t-test is denoted by *. A log2FC>2.0 is denoted by **.

## Supplemental tables

Supplemental table 1 - scRNAseq objects used in this study

Supplemental table 2 - Zebra Finch gene annotations used in this study, with summary statistics Supplemental table 3 - Cell barcodes of the RU HH28 Zebra Finch gonad datasets Supplemental table 4 - Cell Types Markers for CT Inference

Supplemental table 5 - GRC gene paralog Candidates Supplemental table 6 - zGC DEGs

Supplemental table 7 - Statistics for GRC Module Scores

Supplemental table 8 - RU Male zGC DEGs Supplemental table 9 - RU Female zGC DEGs Supplemental table 10 - Male vs Female zGC1 DEGs Supplemental table 11 - Male vs Female zGC2 DEGs

Supplemental table 12 - Cell barcodes and metadata assessed from Jung et al., 2021 (SNU) Supplemental table 13 - SNU zGC DEGs

Supplemental table 14 - Chicken Barcodes Supplemental table 15 - Chicken Genes Supplemental table 16 - cGC vs. cSomatic DEGs Supplemental table 17 - Male vs. female cGC DEGs

Supplemental table 18 - HH28 chicken and zebra finch GC barcodes Supplemental table 19 - HH28 GC Venn Diagram DEG designations Supplemental table 20 - DE GO Terms

Supplemental table 21 - GO Term PC Variance Supplemental table 22 - GO Term PC Loadings

Supplemental table 23 - Estermann et al., 2020 (MU) cell barcodes Supplemental table 24 - MU cGC average gene expression from each dataset Supplemental table 25 - Female E10.5 cGC1 vs. cGC2 DEGs

Supplemental table 26 - E10.5 cGC vs. cSomatic DEGs Supplemental table 27 - MU E10.5 Venn Diagram DEG designations Supplemental table 28 - Male vs. Female E10.5 cGC DEGs

Supplemental table 29 - Conserved zebra finch and chicken GC1 vs. GC2 DEGs Supplemental table 30 - MU reference-query mapping of RU HH28 zGCs Supplemental table 31 - Statistics for MU reference-query mapping Supplemental table 32 - MU reference-query mapping of SNU HH28 zGCs Supplemental table 33 - MU reference-query mapping of RU HH28 cGCs

Supplemental table 34 - Primers used in this study

